# Paradoxical neuronal hyperexcitability in a mouse model of mitochondrial pyruvate import deficiency

**DOI:** 10.1101/2020.12.22.423903

**Authors:** Andres De la Rossa, Marine H. Laporte, Simone Astori, Thomas Marissal, Sylvie Montessuit, Preethi Sheshadri, Eva Ramos-Fernández, Pablo Mendez, Abbas Khani, Charles Quairiaux, Eric Taylor, Jared Rutter, José Manuel Nunes, Alan Carleton, Michael R. Duchen, Carmen Sandi, Jean-Claude Martinou

## Abstract

Neuronal excitation imposes a high demand of ATP in neurons. Most of the ATP derives primarily from pyruvate-mediated oxidative phosphorylation, a process that relies on import of pyruvate into mitochondria occuring exclusively via the mitochondrial pyruvate carrier (MPC). To investigate whether deficient oxidative phosphorylation impacts neuron excitability, we generated a mouse strain carrying a conditional deletion of MPC1, an essential subunit of the mitochondrial pyruvate carrier, specifically in adult glutamatergic neurons. We found that, despite decreased levels of oxidative phosphorylation in these excitatory neurons, mice were normal at rest. Paradoxically, in response to mild inhibition of GABA mediated synaptic activity, they rapidly developed severe seizures and died, whereas under similar conditions the behaviour of control mice remained unchanged. We show that neurons with a deficient MPC are intrinsically hyperexcitable as a consequence of impaired calcium homeostasis, which reduces M-type potassium channel activity. Provision of ketone bodies restores energy status, calcium homeostasis and M-channel activity and attenuates seizures in animals fed a ketogenic diet. Our results provide an explanation for the paradoxical seizures that frequently accompany a large number of neuropathologies, including cerebral ischemia and diverse mitochondriopathies, in which neurons experience an energy deficit.

**One Sentence Summary:** Decreased OXPHOS and Ca^2+^-mediated neuronal hyperexcitability lead to seizure in a mouse model of mitochondrial pyruvate import deficiency.

## Introduction

The brain is by far the main consumer of glucose and oxygen in the body, with pre- and post-synaptic mechanisms being the primary sites of ATP consumption(*1–3*). Most of the ATP in neurons is produced in mitochondria through pyruvate-mediated oxidative phosphorylation (OXPHOS), even though aerobic glycolysis, or the so-called Warburg effect, can generate sufficient ATP to sustain several neuronal functions, including neuronal firing(*3–7*). In neurons, pyruvate is produced either by glycolysis or through the action of lactate dehydrogenase, which mainly uses lactate derived from astrocytes(*8*). Whatever the source, pyruvate transport into mitochondria provides fuel for the tricarboxylic acid (TCA) cycle and boosts ATP production by OXPHOS.

Entry of pyruvate into mitochondria is totally dependent on the mitochondrial pyruvate carrier (MPC), a heterodimer composed of two subunits, MPC1 and MPC2 inserted into the inner mitochondrial membrane(*9, 10*). Deletion of MPC1 or MPC2 is sufficient to inactivate the carrier activity, and in the mouse causes embryonic lethality at E12(*11, 12*). Interestingly, providing ketone bodies, which directly feed the TCA cycle with acetyl-CoA and boost OXPHOS, to the pregnant females allowed the embryos to survive until birth(*12*). Besides energy production, glucose oxidation via the TCA cycle is also required for the synthesis of essential molecules, including the neurotransmitters glutamate and γ-aminobutyric acid (GABA). Therefore ATP production and neurotransmitter release are tightly linked to glucose and pyruvate metabolism. Accordingly, genetic pathologies linked to impaired glucose or pyruvate oxidation, such as mutations in the glucose transporter 1 (GLUT1)(*13*), pyruvate dehydrogenase (PDH)(*13*), MPC(*14, 15*), or complexes of the respiratory chain(*16*), result in severe synaptic dysfunction(*17*). Not surprisingly, these diseases are frequently associated with brain hypoactivity, although paradoxically they are often accompanied by neuronal hyperexcitability and behavioural seizures of varying severity. This raises the question of how these paroxysmal, ATP consuming events can occur in patients despite a global brain energy deficit.

A likely explanation for the hyperexcitable phenotype is that seizures are due to an imbalance between inhibitory (mainly GABAergic) and excitatory (mainly glutamatergic) neuronal activity. While it is understandable that GABAergic neurons may release less inhibitory GABA in the pathologies mentioned above, it remains unclear how excitatory neurons with limited ATP production capacity can display hyperactivity leading to paroxystic seizures within the context of low GABAergic neuronal activity.

To test whether a change in neuronal metabolism would impact neuronal activity, we inactivated the MPC in adult mice, specifically in CamKIIα-expressing neurons (i.e. excitatory neurons) to reduce their OXPHOS capacity, and we analysed their electrical activity at rest and upon pharmacological inhibition of GABAergic transmission. This was achieved through genetic ablation of the MPC1 gene in adult mice using a tamoxifen-inducible system. Furthermore, by using CamKIIα-Cre mice we were able to target the MPC1 deletion and study the impact of decreased pyruvate oxidation specifically in glutamatergic neurons. We found that, under resting conditions, mice lacking MPC1 in these excitatory neurons were indistinguishable from control mice in their general exploratory, social and stress-coping behaviors. In response to inhibition of GABA mediated synaptic activity they developed far more severe seizures than controls. We found that this phenotype was due to an intrinsic membrane hyperexcitability of MPC1-deficient glutamatergic neurons, which resulted from a calcium-mediated decrease in M-type K^+^ channel activity. Strikingly, the hyperexcitability phenotype was reversed when the animals were maintained on a ketogenic diet.

## Results

### MPC-deficient cortical neurons display decreased pyruvate-mediated oxidative phosphorylation in vitro

To assess the role of the mitochondrial pyruvate carrier (MPC) in neuronal OXPHOS, we first used primary cultures of cortical neurons largely depleted of astrocytes (Supplementary figure 1a) and either RNA interference or pharmacological reagents to downregulate their MPC activity. To this end, two different shRNAs targeting MPC1 and three different pharmacological inhibitors of the carrier were used. Expression of either of the two shRNAs produced a significant reduction in MPC1 and MPC2 protein levels (the latter being unstable in the absence of MPC1) (Supplementary figure 1b, c). Both genetic and pharmacological impairment of MPC activity resulted in decreased pyruvate-driven basal and maximal oxygen consumption rates (OCR) (Figure 1a, Supplementary figure 1d) and decreased mitochondrial ATP production (Figure 1b), which is consistent with previously published results(*18*). Furthermore, mitochondrial membrane potential, measured using mitotracker and TMRE was significantly reduced in MPC-deficient neurons (Figure 1c, f). This was associated with an increased extracellular acidification rate (Supplementary figure 1e) and increased glucose uptake, which was measured using the 2-NBDG import assay (Supplementary figure 1f), two hallmarks of aerobic glycolysis.

**Figure 1.**
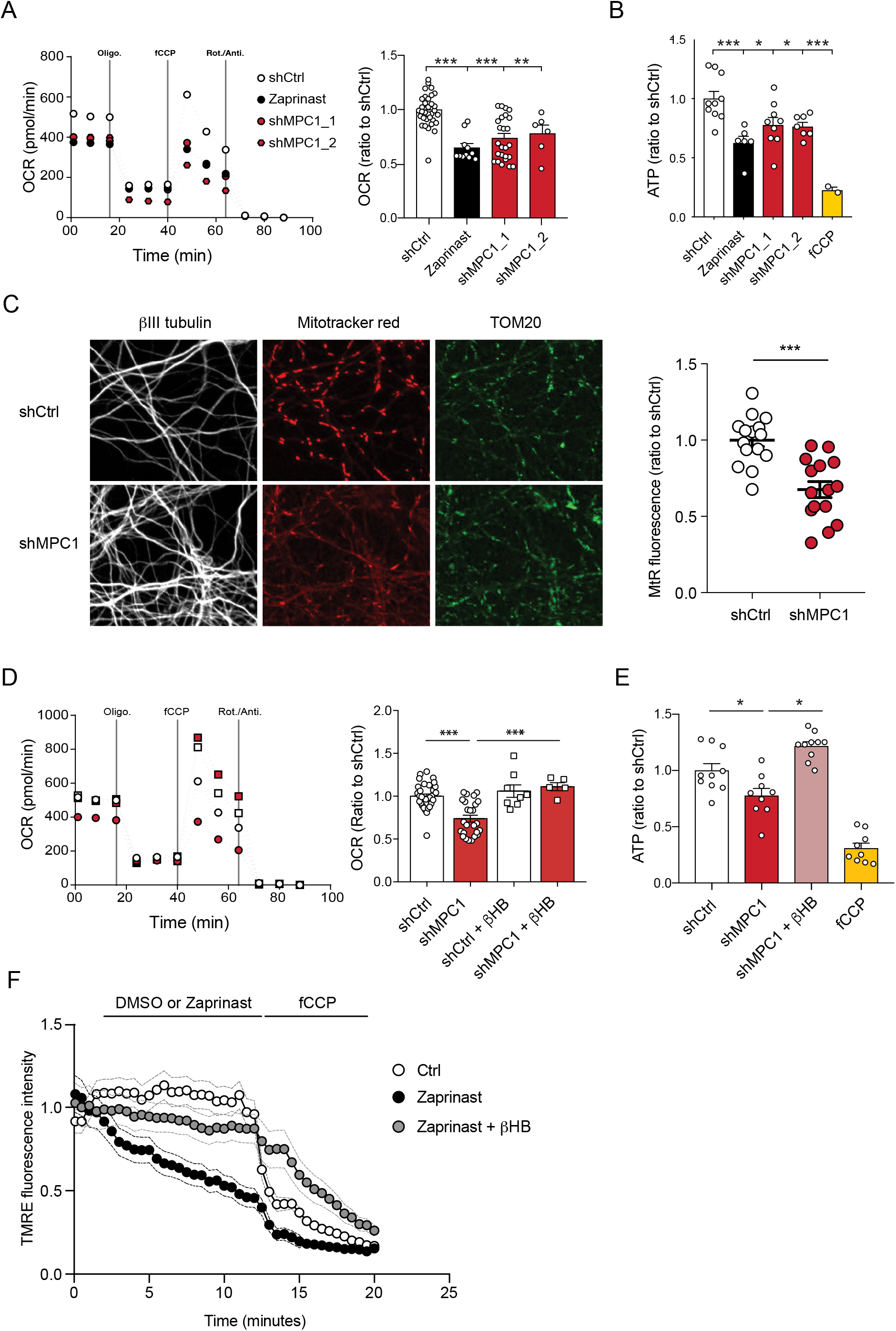
MPC-deficient neurons display defects in mitochondrial respiration and membrane potential. **A)** Profile and quantification of oxygen consumption rates (OCR) cortical neurons expressing either shCtrl, or shMPC1_1 and shMPC1_2 for 7 days, or in the presence of Zaprinast (5μM, 1 hour). Data were obtained using the Seahorse XF analyzer. Assays were performed in the presence of pyruvate (5 mM) and glucose (5 mM) as carbon sources. Quantification of basal OCR is expressed as ratio of ShCtrl. N=10,7,9,7 and 2 independent experiments. N=33,11,25 and 6 independent experiments. One-way ANOVA+Tukey’s post-hoc test (shCtrl vs Zaprinast p=0.0001, shCtrl vs shMPC1_1 p=0.0001, shCtrl vs shMPC1_2 p=0.0013). **B)** ATP content in MPC-deficient cortical neurons treated with either shCtrl, Zaprinast or shMPC1_1 and shMPC1_2. fCCP (4 μM) treatment reveals the non-mitochondrial ATP. N=10,7,9,7 and 2 independent experiments. One-way ANOVA+Tukey’s post-hoc test (shCtrl vs Zaprinast p=0.0006, shCtrl vs shMPC1_1 p=0.0223, shCtrl vs shMPC1_2 p=0.0242, shCtrl vs fCCP p=0.0001). **C)** Mitochondrial membrane potential of MPC-deficient cortical neurons. Neurons were incubated with Mitotracker red (MtR) (1 μM) prior fixation, immunostained for βIII tubulin (neuron) and TOM20 (mitochondria). Quantification of Mitotracker red fluorescence in each βIII tubulin-positive cell (red) was reported to TOM20 signal (green). N=15 neurons from 3 independent experiments. Unpaired t test (shCtrl vs shMPC1 p=0.0001). **D)** Profile and quantification of oxygen consumption rates (OCR) in cortical neurons expressing shCtrl or shMPC1_1 for 7 days. Data were obtained using the Seahorse XF analyzer. Assays were performed in the presence of pyruvate (5 mM) and glucose (5 mM) as carbon sources + 10 mM βHB when indicated. Quantification of basal OCR is expressed as ratio of control condition shCtrl. N=12 independent experiment. One-way ANOVA+Holm Sidak’s post-hoc test (shCtrl vs shMPC1 p=0.0001, shMPC1 vs shMPC1+βHB p=0.0002). **E)** ATP content in MPC-deficient cortical neurons treated with shCtrl or shMPC1 in presence or absence of 10 mM βHB. fCCP (4 μM) treatment reveals the non-mitochondrial ATP. N=10, 9, 10, 9 independent experiments. One-way ANOVA+Holm Sidak’s post-hoc test (shCtrl vs shMPC1 p=0.0145, shMPC1 vs shMPC1+βHB p=0.0143, shCtrl vs fCCP p=0.0001). **F)** Cortical neurons were incubated with TMRE (50 nM) +/- βHB (10 mM) for 30 min and recorded by live microscopy. Neurons were incubated with DMSO or Zaprinast (5 μM) 2.5 min after the beginning of the acquisition and recorded for 5 min prior fCCP injection. N=15 independent experiments. One-way ANOVA+Holm Sidak’s post-hoc test (shCtrl vs Zaprinast p=0.0028, Zaprinast vs Zaprinast+βHB p=0.0008).

We have previously reported that ketone bodies can restore normal OXPHOS in MPC-deficient murine embryonic fibroblasts(*12*). Consistent with this, we found that addition of the ketone body β-hydroxybutyrate (βHB) to the culture medium rescued all observed defective functionalities in MPC-deficient neurons, including oxygen consumption, ATP production, membrane potential (Figure 1d-f) and both extracellular acidification rate and glucose uptake (Supplementary figure 1g, h). Thus, we concluded that MPC-deficient neurons display low pyruvate-mediated oxidative phosphorylation and high aerobic glycolysis, both overcomed with βHB.

### Generation of mice with inducible MPC1 gene deletion in adult glutamatergic neurons

Based on the results described above, and because neural excitation requires massive levels of ATP, we hypothesized that loss of MPC activity would reduce excitability especially in glutamatergic neurons that are high energy consumers(*19*). To test this hypothesis, we generated a mouse strain with an inducible deletion of the MPC1 gene, specifically in the Ca^2+^-calmodulin kinase IIα (CamKIIα)-expressing neurons, found predominantly in the hippocampus and cortex(*20*). We crossed MPC1^flox+/flox+^ mice with the commercially available CamKIIα-CreERT2 mice (Figure 2a). Induction of Cre activity by injection of Tamoxifen for 5 consecutive days resulted in deletion of MPC1 FLOXed alleles in the CamKIIα-expressing adult neurons (Figure 2a). Hereafter, we refer to these mice as neuro-MPC1-KO. *In situ* immunofluorescence analyses showed a decrease in neuronal MPC1 immunostaining in various layers of the cortex of neuro-MPC1-KO mice (Figure 2b). Western blot analysis of whole cortex, synaptosomes and mitochondria showed a significant decrease of both MPC1 and MPC2 in neuro-MPC1-KO mice compared to neuro-MPC1-WT mice (Figure 2c). Consistent with the results obtained with cultured neurons, we found that synaptosomes prepared from the cortex of neuro-MPC1-KO mice displayed lower oxygen consumption and imported higher amounts of glucose compared to synaptosomes from neuro-MPC1-WT mice (Supplementary figure 2a, b). Importantly, the lack of MPC1 did not affect the neuronal cell survival quantified either by counting the total number of cells, or by the number of apoptotic (TUNEL positive) cells (Supplementary figure 2c, d). At adulthood, both genotypes displayed similar body weight and lean mass composition (Supplementary figure 2e, f). At the behavioral level, adult neuro-MPC1-KO mice showed a tendency toward lower anxiety-like behaviors, but no difference in general locomotion, sociability or stress-coping behaviors (Supplementary figure 2g-j).

**Figure 2.**
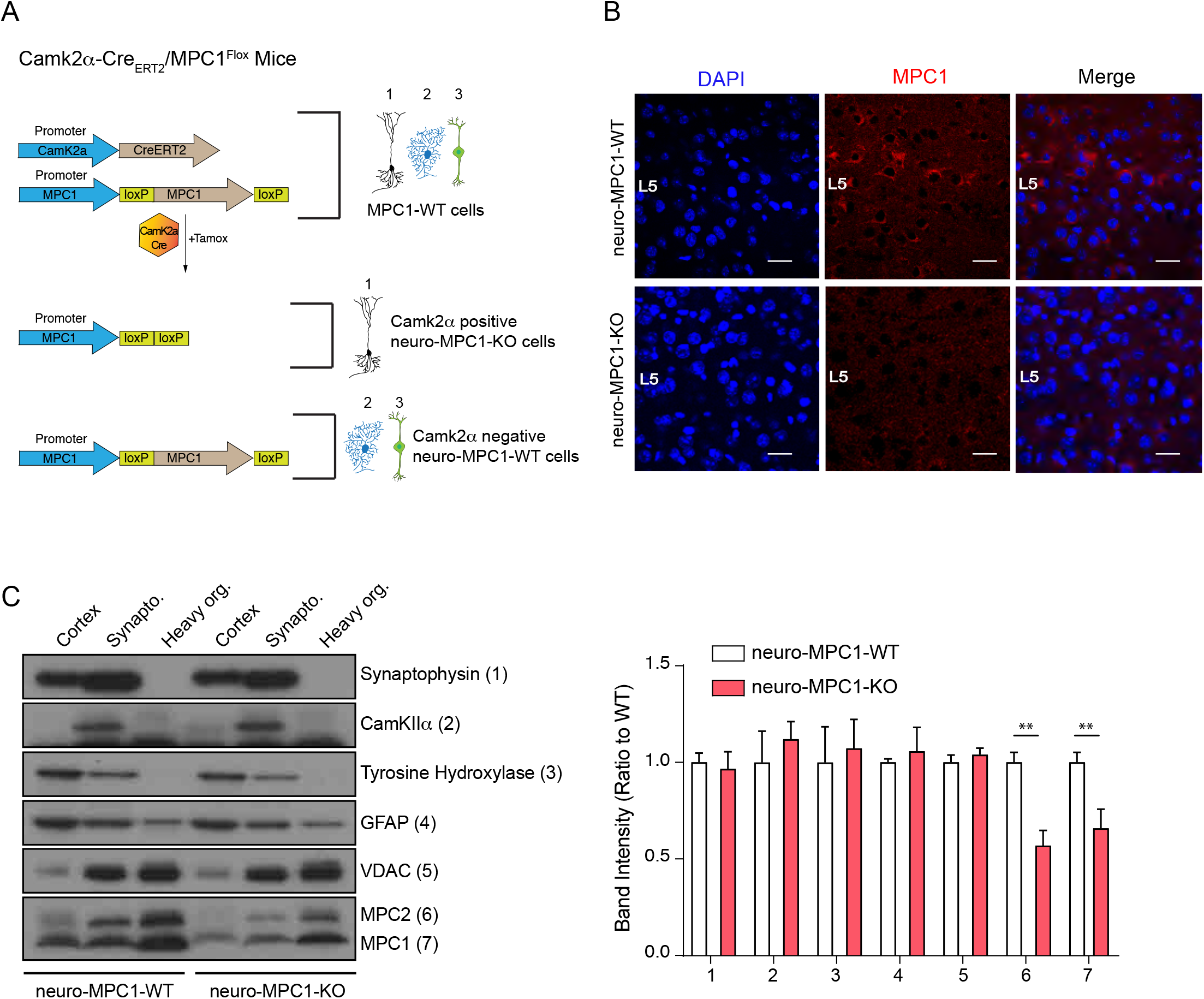
Generation of mice with an inducible deletion of the MPC1 gene in adult glutamatergic neurons. **A)** Strategies used to generate CamKIIα-Cre_ERT2_/MPC1^Flox^ mice. Upon Tamoxifen injection, expression of the Cre recombinase in CamKIIα glutamatergic neurons drives deletion of the MPC1 gene. These mice are referred to as neuro-MPC1-KO or neuro-MPC1-WT when they are CamCre-(1. Glutamatergic neuron; 2. Astrocytes; 3. Inhibitory neuron). **B)** Immunostaining of MPC1 (red) in cortical sections from neuro-MPC1-WT and neuro-MPC1-KO mice (scale bar: 100 μm). **C)** Western blot analysis of whole cortex, synaptosome lysates and heavy organelles (mainly mitochondria), obtained from brains of neuro-MPC1-WT and neuro-MPC1-KO mice using neuronal (Synaptophysin, tyrosine hydroxylase, CamKIIα) and astroglial markers (GFAP) as well as mitochondrial markers (MPC1, MPC2 and VDAC). Note that synaptosomes are enriched for CamKIIα, a marker of excitatory neurons. Quantification (right panel) shows that except for MPC1 and MPC2, the content of these markers is similar in WT and KO preparations. N=6 independent neuro-MPC1-WT and neuro-MPC1-KO mice. Mann-Whitney test ((6) neuro-MPC1-WT vs neuro-MPC1-KO p=0.0286, (7) neuro-MPC1-WT vs neuro-MPC1-KO p=0.0152).

These data indicate that, under resting conditions, the excitatory neurons in most adult mice have the ability to bypass the MPC to meet their metabolic demands.

### Neuro-MPC1-KO mice are highly sensitive to pro-convulsant drugs and develop acute epileptic-like seizures

The output activity of a neuron results from the balance between the excitatory and the inhibitory inputs it receives. Perturbation of this delicate balance can lead to severe seizures as a result of exacerbated, uncontrolled neuronal firing. To test whether OXPHOS-deficient excitatory neurons could sustain intense neuronal firing, we challenged neuro-MPC1-KO adult mice, with either pentylenetetrazole (PTZ), a GABA receptor antagonist, or kainic acid, an activator of glutamate receptors. We used the PTZ kindling protocol described previously(*21*), in which a sub-convulsant dose (35mg/kg) of PTZ is injected intraperitoneally (ip) once every two days on a period of 15 days (Figure 3a). Phenotypic scoring after each PTZ injection in neuro-MPC1-WT mice showed a progressive sensitization (kindling) starting with hypoactivity after the first injection (scored as 1); a few brief and transient muscle contractions (jerks, scored as 2) or appearance of tail rigidity (Straub’s tail, scored as 3) following the second or third injection; and convulsive status epilepticus (scored as 6) after the 6^th^ or 7^th^ injection (Figure 3a, b). In contrast, all neuro-MPC1-KO mice developed severe, prolonged seizures (score 6) within 10 min of the first PTZ injection (Figure 3b) and all died during seizures within the next three PTZ injections (Supplementary figure 3a). When mice were injected with 20 mg/kg kainic acid, a similar hypersensitivity (score 6) was observed in neuro-MPC1-KO mice indicating that this sensitivity is not restricted to PTZ (Supplementary figure 3b).

**Figure 3.**
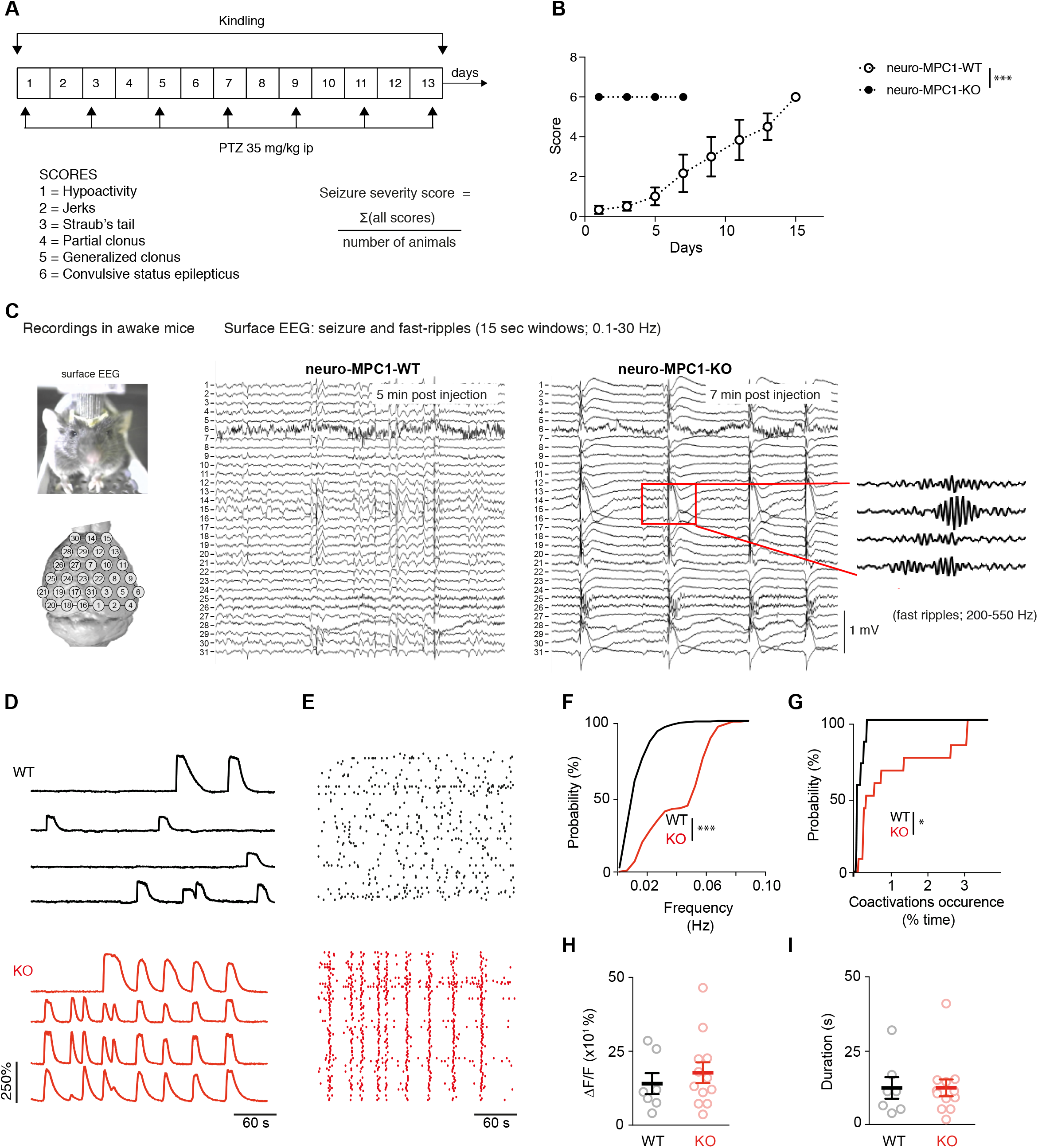
Neuro-MPC1-KO mice are highly sensitive to pro-convulsant drugs and develop acute epileptic-like seizures. **A)** Schematic description of the PTZ kindling protocol. **B)** Seizure severity scores reflecting the different clinical symptoms as indicated, obtained for neuro-MPC1-WT or neuro-MPC1-KO. N=8 independent neuro-MPC1-WT and neuro-MPC1-KO mice. Two way ANOVA (F(7,70)=19, p=0.0001). **C)** Illustration of the recording setups in awake mice indicating the position of surface EEG electrodes and representative example of a seizure recorded in a neuro-MPC1-KO mouse after injection of 35mg/kg PTZ during surface EEG recordings. The inset shows an example of fast ripples generated during an ictal epileptic discharge. **D-I)** GCaMP6S calcium imaging of the CA1 area from hippocampal slices in the presence of Carbachol (50μM) and PTZ (2mM). Slices were prepared from WT animals (top, black) or from KO animals with no pre-treatment (bottom, red). **D)** Ca^2+^ sweeps recorded in four representative GCaMP6S-expressing neurons. **E)** Raster plots of Ca^2+^ transient onsets extracted from all recorded neurons in a given slice. **F)** Cumulative distribution of the frequency of the calcium events in all the recorded neurons. N=7, 12 independent experiments. Kolmogorov-Smirnov test (WT vs KO p=0.0001). **G)** Cumulative distribution of the occurrence of neuronal co-activations exceeding chance levels as a function of time N=7, 12 independent experiments. Kolmogorov-Smirnov test (WT vs KO p=0.0344). Amplitude (**H**), and duration (**I**) of the calcium events recorded in all neurons of the hippocampus. N=7, 12 independent experiments. Mann-Whitney test (Amplitude: WT vs KO p=0.5918; Duration: WT vs KO p=0.9182).

In a parallel series of experiments, and in order to assess the specificity of our results to excitatory neurons, we investigated the effects of PTZ in mice in which MPC1 was deleted in adult astrocytes (hereafter termed astro-MPC1-KO mice) (Supplementary figure 3c-e). In contrast to neuro-MPC1-KO mice, astro-MPC1-KO mice showed the same response as control animals following PTZ injection (Supplementary figure 3f, g), indicating that the phenotype observed in neuro-MPC1-KO mice is linked to the deletion of MPC1 in excitatory neurons.

To characterize the seizure symptoms in more detail, we recorded the electrical activity in the brains of neuro-MPC1-WT and neuro-MPC1-KO mice by electroencephalogram (EEG) following a single injection of PTZ (Figure 3c). In neuro-MPC1-KO mice, rhythmic EEG patterns emerged within 5-10 minutes after PTZ injection, invading all electrodes (Figure 3c). These electrical patterns coincided with the occurrence of behavioural manifestations of seizures, i.e. tonic-clonic movements. Rapidly thereafter, large spike and wave discharges developed, again invading all surface electrodes and coinciding with numerous fast ripples (Figure 3c, inset). Such EEG patterns are characteristic of seizure episodes in humans and were not observed in the PTZ-injected neuro-MPC1-WT mice. These data indicate that neuro-MPC1-KO mice develop an epilepsy-like phenotype following administration of a single sub-convulsant dose of PTZ.

We also tested whether we could reproduce the seizure phenotype using hippocampal organotypic cultures from CamKIIα-CreERT2^+^-MPC1^Flox+/Flox+^ mice exposed to PTZ, combined with calcium imaging. Individual neurons in hippocampal slices from both WT and KO mice exhibited spontaneous calcium activity throughout the duration of the recordings (Figure 3d, e and videos 1 and 2) although, interestingly, the frequency of calcium events, as well as the number of co-activation events (i.e. neuronal synchronizations above chance levels) generated in MPC1-deficient neurons were significantly higher than those generated in MPC1-WT neurons (Figure 3f, g and videos 1 and 2). In contrast, neither the amplitude nor the duration of the discharges was modified (Figure 3h, i). These results suggest that neuro-MPC1-KO neurons are more active and are more often recruited into synchronized patterns associated with the epileptic activity.

### Inhibition of PTZ-induced seizures in neuro-MPC1-KO mice by the ketogenic diet

The ketogenic diet (KD) has been reported to decrease seizures in patients with pharmacologically refractory epilepsy(*22*). Ketone bodies, mainly generated by the liver during fasting and hypoglycaemia, are used by neurons to provide the TCA cycle with acetyl-CoA, normally provided by pyruvate dehydrogenase-mediated oxidation of pyruvate. Thus, ketone bodies ensure that oxidative phosphorylation and ATP production is maintained in neurons in conditions of glucose starvation. We tested whether a ketogenic diet could prevent PTZ-induced seizures in neuro-MPC1-KO mice. As previously reported(*23*), we found that the KD produces a decrease in glycaemia and an increase in the blood level of 3-β-hydroxy-butyrate (βHB), one of the three major ketone bodies generated by the liver (Supplementary figure 4a, b). In addition, we found that mice fed on the KD for one week were completely resistant to PTZ injection (Figure 4a). Supplementing the drinking water with 1% βHB was sufficient to prevent PTZ-induced seizures (Figure 4b). Similarly, ip administration of βHB 15 minutes before PTZ injection, or starvation overnight, both of which conditions led to increased βHB blood levels (Supplementary figure 4c-e), significantly reduced the PTZ-induced clinical score of neuro-MPC1-KO mice (Figure 4b, c). These results indicate that the phenotype displayed by the neuro-MPC1-KO mice is mainly metabolic in origin and is unlikely to be the consequence of neuronal network remodelling.

**Figure 4.**
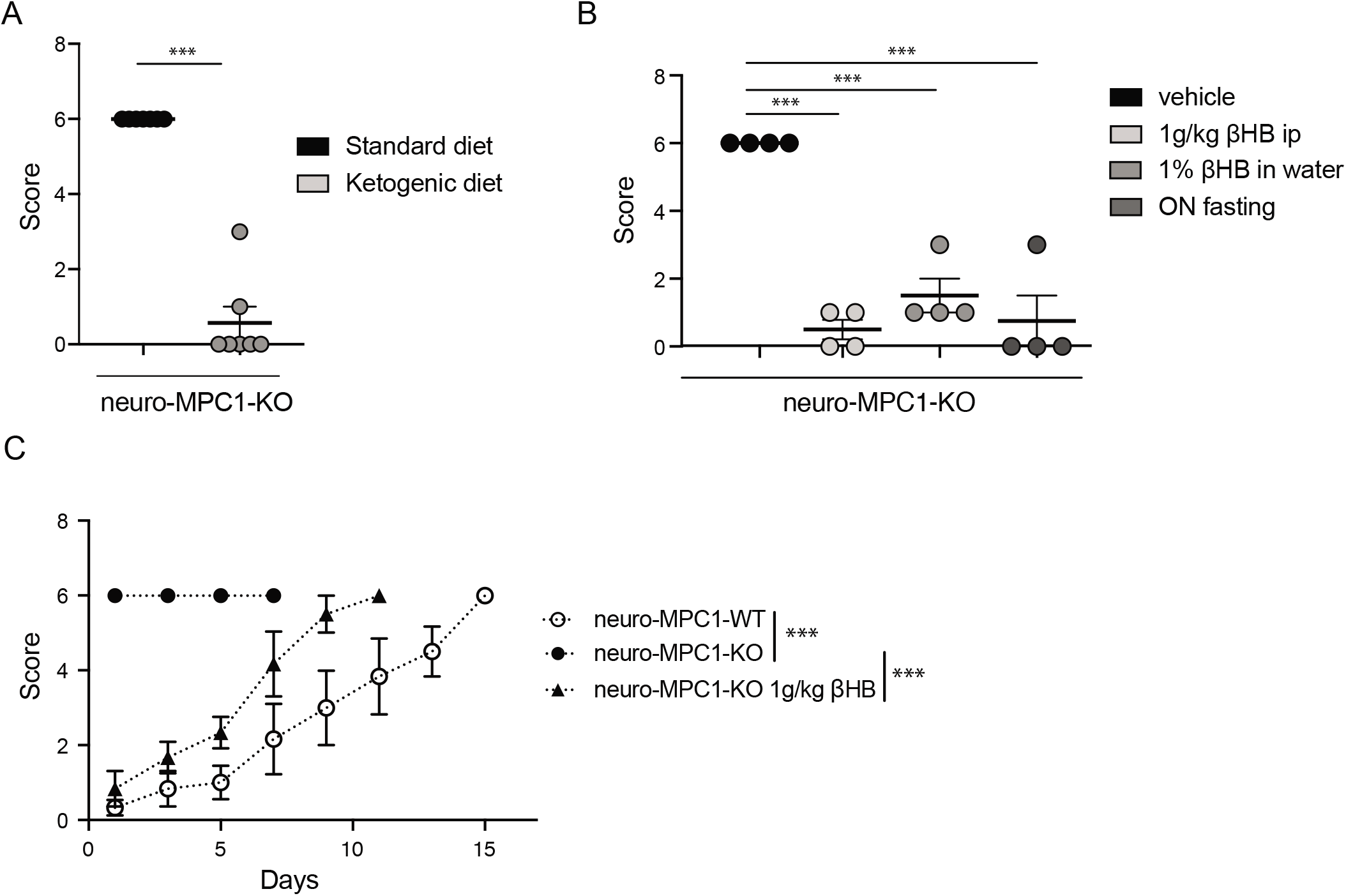
Ketogenic diet prevents the epileptic phenotype of neuro-MPC1-KO mice. **A)** Effect of the ketogenic diet (KD) on PTZ-induced seizure. All neuro-MPC1-KO mice were maintained on the Standard (SD) or ketogenic (KD) diet for 7 days prior to challenge with a single dose of PTZ. Clinical scores were assessed directly following injection. N=7 independent neuro-MPC1-KO mice. Mann-Whitney (neuro-MPC1-KO SD vs neuro-MPC1-KO KD p=0.0008). **B)** Effects 1% βHB in the drinking water for 7 days, overnight fasting or ip injection of βHB 15 min before administration of PTZ into neuro-MPC1-KO mice. N=4 independent neuro-MPC1-KO mice. One-way ANOVA+Holm Sidak’s post-hoc test (Vehicle vs all conditions p=0.0001). **C)** Effect of βHB on PTZ-induced seizure: mice were injected ip with 1g/kg βHB, 15 minutes before each PTZ injection and scored for clinical symptoms. N=6 independent mice. Two-way ANOVA+Holm Sidak’s post-hoc test (F(10, 75)=8, Neuro-MPC1-WT vs neuro-MPC1-KO, neuro-MPC1-KO vs neuro-MPC1-KO + βHB p=0.0001).

### MPC1-deficient neurons display intrinsic hyperexcitability, which is prevented by ketone bodies

To investigate the cellular mechanisms that might mediate the sensitivity of neuro-MPC1-KO mice to pro-convulsant drugs, we examined the electrophysiological properties of MPC1-deficient neurons. To this end, we performed whole-cell patch clamp recordings in acute hippocampal slices from neuro-MPC1-KO mice and their neuro-MPC1-WT littermates. CA1 pyramidal cells from neuro-MPC1-KO mice exhibited higher discharge frequency compared to neurons from neuro-MPC1-WT mice when firing was elicited by somatic injections of current ramps of increasing amplitude (Figure 5a, b). Neurons from neuro-MPC1-KO mice required less current injection (rheobase, Figure 5c) to reach the firing threshold, which was more hyperpolarized when compared to neuro-MPC1-WT cells (Figure 5d). Similarly, MPC1-KO neurons displayed higher firing when depolarization was induced with squared current pulses (Supplementary figure 5a, b).

**Figure 5.**
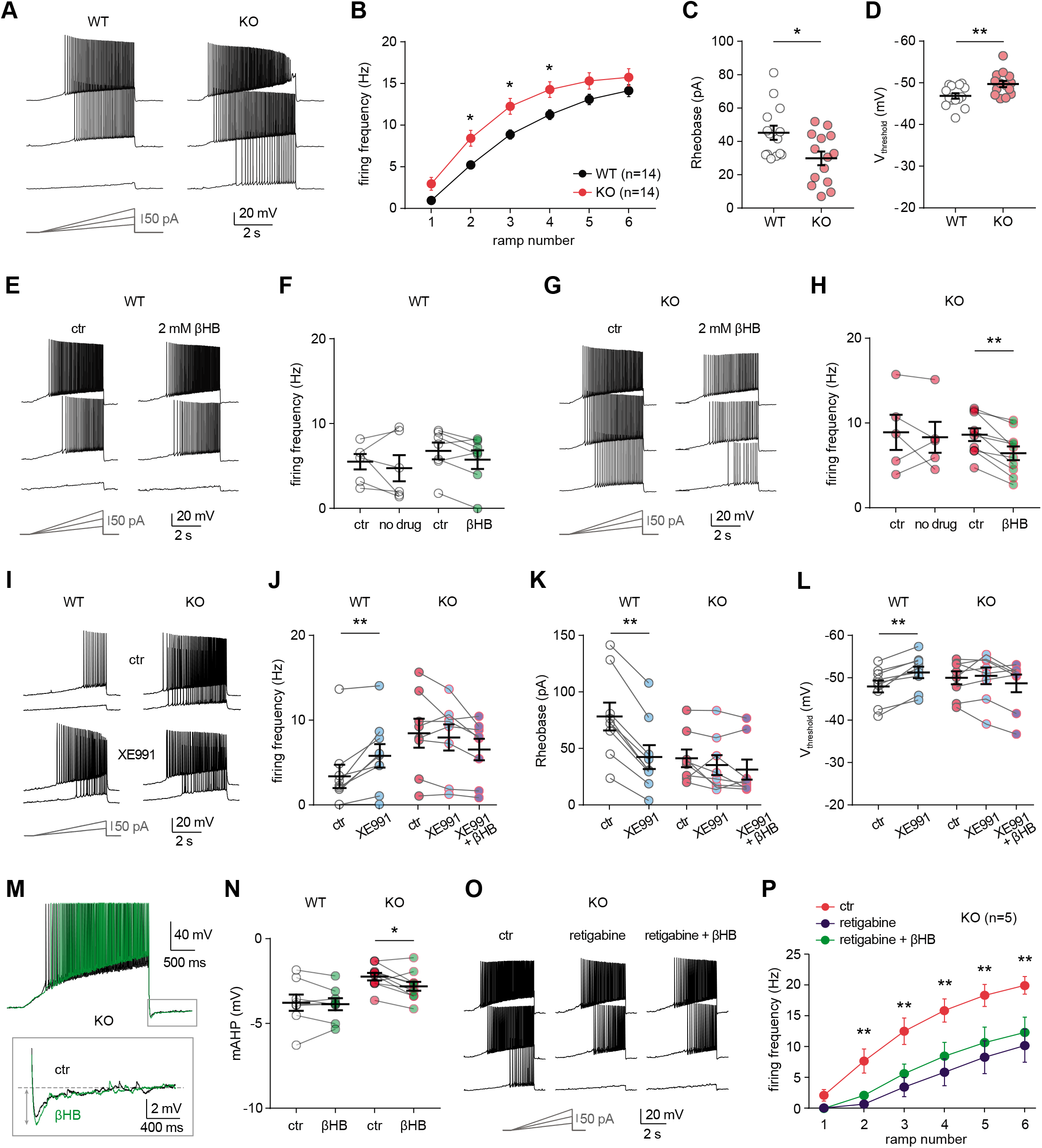
MPC1 deletion increases intrinsic excitability in CA1 pyramidal cells. **A)** Example voltage responses elicited in CA1 pyramidal cells from wild-type (WT) and MPC1-CamKII-KO (KO) by injection of current ramps (protocol at the bottom, only three of six ramps displayed). **B)** Frequency-current (F-I) relationship of action potential discharges, indicating higher spiking frequency in KO cells (Two-way ANOVA, F(1, 156) 33.43, p<0.0001). **C)** The rheobase was reduced in KO cells (Mann-Whitney test, U=53.5, p=0.0406). **D)** KO cells exhibited more hyperpolarized threshold potential (unpaired t test, t=2.856, p=0.0084). **E)** Example traces showing lack of changes in WT cell firing after bath application of the ketone body β-hydroxybutyrate (βHB, 2 mM, >20 min exposure). **F)** Average firing frequency elicited by the 3^rd^ ramp of current injections in WT cells in control condition (5 min after whole-cell establishment) and after 20 min of either no drug exposure or βHB application (no drug: paired t test, t=0.664, p=0.5362; βHB: paired t test, t=2.1, p=0.0804). **G, H)** Example traces and summary graphs indicating significant reduction in KO cell firing after βHB application (no drug: paired t test, t=0.4691, p=0.6634; βHB: paired t test, t=5.339, p=0.0005). **I)** Example traces of cell firing in control and after bath application of XE991 (10 μM) in WT and KO neurons. **J)** Average firing frequency elicited by the 2^nd^ ramp of current injections in control and after XE991 application, indicating increased excitability in WT cells (paired t test, t=3.735, p=0.0057). XE991 was ineffective in KO cells, in which subsequent application of βHB also failed to modulate excitability (One-way ANOVA, F(1.69, 11.89)=4.76, p=0.0347, Holm-Sidak’s multiple comparison p>0.05). **K)** XE991 significantly reduced the rheobase of WT cells (paired t test, t=11, p<0.001), but not of KO cells, in which subsequent βHB application was also ineffective (One-way ANOVA, F(1.785, 12.5)=2.99, p=0.091). **L)** XE991 induced a shift in the threshold potential of WT cells (paired t test, t=6.001, p=0.0003), but did not affect KO cells, in which subsequent βHB application was also ineffective (One-way ANOVA, F(1.812, 12.68)=1.78, p=0.209). **M)** Example traces of KO cell firing elicited by a current ramp (300 pA max amplitude, APs are trimmed) in control and after βHB exposure, with expanded portion at the bottom indicating mAHP measurement. **N)** Summary graph of mAHP values in WT and KO cells in control and after βHB exposure, indicating significant increase in KO (unpaired t test, t=2.89, p=0.0179). **O)** Example traces of KO cell firing before and after application of retigabine (10 μM), and subsequent βHB superfusion (2 mM). **P)** F-I relationships in KO cells, indicating reduced spiking frequency after retigabine application, with no additional effect of βHB (Two-way repeated measures ANOVA, F(2, 48)=89.15, p<0.0001).

Next, we asked whether ketone bodies, which as shown in Figure 4 prevent PTZ-induced seizures, could modulate neuronal excitability and restore normal cell discharges in neuro-MPC1-KO mice. For these experiments, we first recorded action potential firing under control conditions, and then perfused the slices with βHB (2 mM, >20 min exposure). As shown in Figure 5, whereas cell firing was unaltered in neuro-MPC1-WT cells (Figure 5e, f), βHB reduced excitability in pyramidal cells from the neuro-MPC1-KO mice (Figure 5g, h). Control experiments showed that cell excitability from both genotypes was unchanged during prolonged recordings (Figure 5f, h), confirming that the change in neuro-MPC1-KO firing was not due to a rundown in cellular excitability caused by, e.g., cell dialysis.

Taken together, these results indicate that ketone bodies reduce the intrinsic hyperexcitability of glutamatergic cells from neuro-MPC1-KO mice, providing a plausible explanation for the protective effect of the KD against PTZ-induced seizures.

### MPC1-deficient neurons display altered M-type potassium channel activation, which is corrected by β-hydroxybutyrate

To gain insight into the mechanisms governing neuronal hyperexcitability, we analysed the cellular passive properties and action potential characteristics of all recordings performed in cells from neuro-MPC1-KO and neuro-MPC1-WT mice (Supplementary figure 5c-k). The reduction in rheobase and the shift in threshold potential induced by MPC1 deletion were accompanied by several changes in passive and active membrane properties governing cell excitability, including a significant increase in the input resistance (Ri) and in the voltage response to a depolarizing current injection (depol_sub_), along with a marginally significant reduction in HCN channel-mediated sag (Supplementary figure 5c-i). The fast afterhyperpolarization (fAHP) accompanying action potentials was not altered, ruling out a major contribution of BK channels (Supplementary figure 5j). However, the medium afterhyperpolarization (mAHP), measured as the negative peak of the voltage deflection at the offset of the depolarizing ramps was significantly reduced in cells from neuro-MPC1-KO mice (Supplementary figure 5k). In CA1 pyramidal cells, mAHP is primarily mediated by the activation KCNQ2/3 (Kv7.2 and Kv7.3) channels, which generate an M-type K^+^ conductance regulating intrinsic excitability and synaptic integration(*24, 25*). Opening of these channels produces an outward potassium current that functions as a ‘brake’ for neurons receiving persistent excitatory input(*26*). Consistently, mutations in KCNQ2/3 genes have been associated with seizures in the mouse(*27*), as well as in patients(*28, 29*), pointing to these channels as interesting targets for anticonvulsant therapy(*30*). To verify whether neuro-MPC1-KO mice display an altered contribution of the M-type K^+^ conductance, we tested the effect of the M-type channel blocker XE991 (10 μM) on CA1 pyramidal cell firing. XE991 led to a significant increase in firing frequency of neuro-MPC1-WT cells, whereas firing of neuro-MPC1-KO cells was not significantly modified (Figure 5i, j). Consistently, XE991 induced a significant reduction in the rheobase and a shift in the threshold potential in neuro-MPC1-WT cells, but had no impact on neuro-MPC1-KO cells (Figure 5k, l), pointing to a limited activity of KCNQ2/3 channels in these neurons. Interestingly, bath application of βHB following KCNQ2/3 channel blockade with XE991 failed to reduce the hyperexcitability of neuro-MPC1-KO mice (Figure 5j-l). We also noticed that, in the absence of XE991, the reduction of intrinsic excitability by βHB in MPC-deficient neurons was accompanied by a significant increase in mAHP (Figure 5m, n), suggesting that βHB may potentiate the recruitment of the M-type K^+^ channels. Moreover, the M-type channel activator retigabine (10 μM) effectively decreased the hyperexcitability of pyramidal cells from neuro-MPC1-KO mice to a level that was no further affected by βHB (Figure 5o, p). This suggests that βHB and retigabine display a similar mechanism of action, which is consistent with recent findings showing that βHB can directly bind to and activate KCNQ2/3 channels(*31*).

We finally tested whether the increased neuronal excitability in neuro-MPC1-KO mice was also accompanied by alterations in glutamatergic transmission. In acute slices, we recorded field potentials in CA1 stratum radiatum elicited by electrical stimulation of the Schaffer collaterals (Supplementary figure 5l). No overt genotype differences were found in the input-output curves of field excitatory postsynaptic potentials (fEPSPs), and the lack of changes in paired-pulse ratio indicated no major alteration in the presynaptic release (Supplementary figure 5m, n).

Altogether, these results indicate that the hyperexcitability of CA1 pyramidal neurons from neuro-MPC1-KO mice is mediated by alterations in intrinsic cell excitability associated with a reduced M-type K^+^ channel activation, with no major changes in excitatory synaptic inputs.

### Alteration of calcium homeostasis in MPC1-deficient neurons

The conductance of KCNQ channels is regulated by phosphatidylinositol-4,5-bisphosphate (PIP2) and calmodulin (CaM)(*32, 33*). In particular, reduction in free CaM in hippocampal neurons decreases M-current density and increases neuronal excitability(*34, 35*). Thus, calcium can trigger loss of interaction of CaM and KCNQ2/3 channels, leading to M-type current suppression(*36*).

We tested whether disruption of calcium homeostasis could be responsible for the deficit in the M-type K^+^ channel activity displayed by MPC1-deficient neurons. We first assess whether calcium homeostasis was perturbed in MPC-deficient cortical neurons in vitro. Using the fluorimetric calcium probes Fura2-AM and the low affinity FuraFF-AM and live cell imaging, we found a significant increase in the peak concentration of cytosolic calcium upon depolarization of both control and MPC-deficient neurons in response to either glutamate (10 μM) or KCl (50 mM) (Figure 6a-c; Supplementary figure 6a-d). However, while the peak of calcium concentration was transient in control neurons, and returned to basal levels, both the magnitude and duration of the calcium elevation were greater in MPC-deficient neurons (Figure 6a, Supplementary figure 6d, Wash). Interestingly, the long lasting increased calcium level in MPC-deficient neurons was abolished by addition of 10 mM βHB to the culture medium 30 min prior to recording (Figure 6a-c). Together these results show that loss of MPC activity leads to a significant increase of cytosolic calcium levels in depolarized neurons.

**Figure 6.**
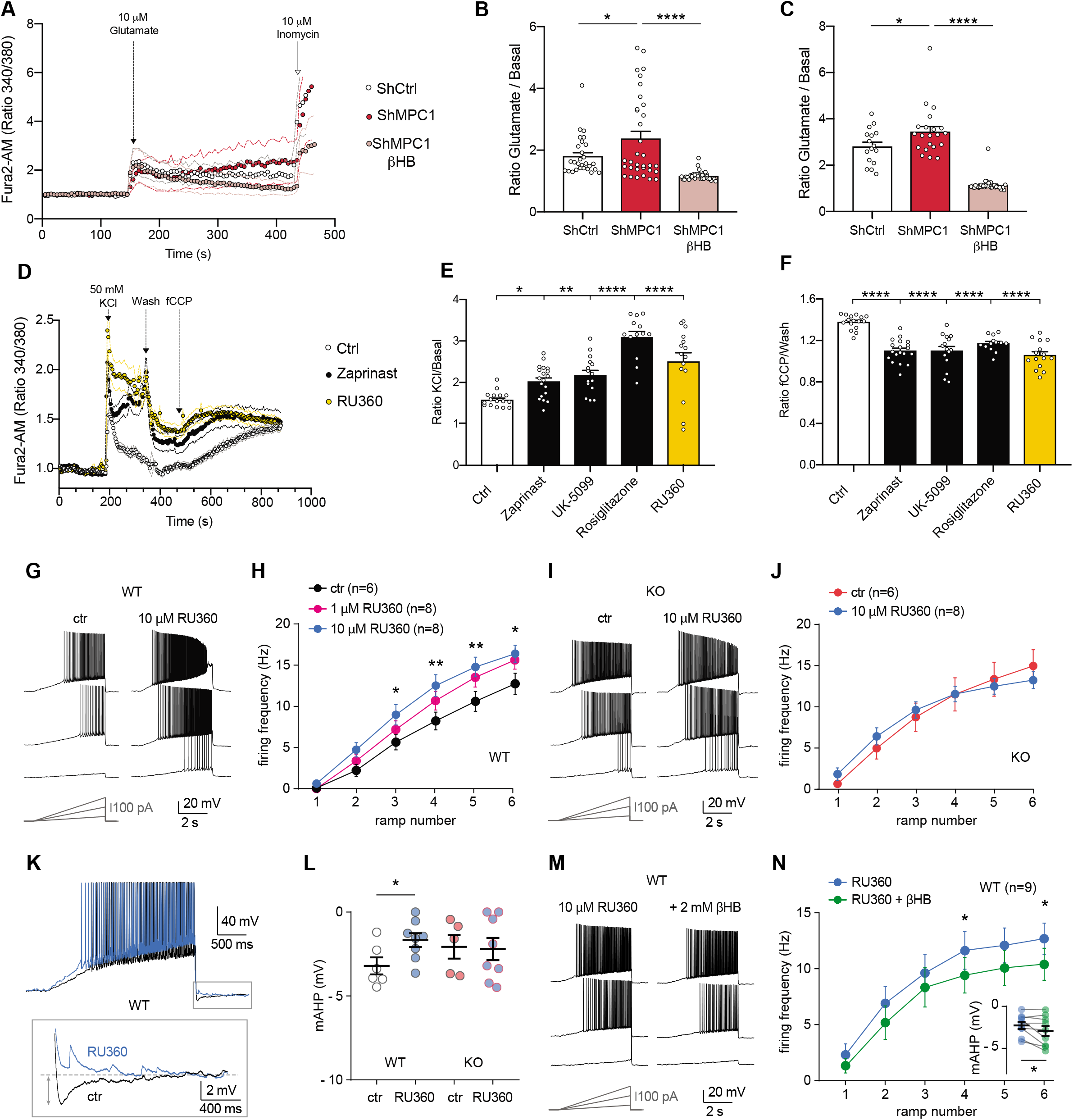
Defect in calcium homeostasis. **A)** Mean fluorescence signal intensity of cortical neurons loaded with furaFF-AM stimulated with 10 μM glutamate (dashed black arrow) prior the addition of ionomycin (red arrow) to reveal the neuronal calcium stock. **B, C)** Graph showing the quantifications of control neurons, MPC-depleted neurons and MPC-depleted neurons+βHB showing an elevated level of cytosolic calcium in MPC-deficient stimulated neurons measured by FuraFF-AM (**B**) or Fura2-AM (**C**). N>15 neurons per condition from 3 independent experiments. One-way ANOVA+Holm Sidak’s post-hoc test ((**B)** shCtrl vs shMPC1 p=0.0286, shMPC1 vs shMPC1+βHB p=0.0001; (**C**) shCtrl vs shMPC1 p=0.0169, shMPC1 vs shMPC1+βHB p=0.0001). **D**) Fluorescence signal intensity of control, MPC-deficient, and RU360-treated cortical neurons permeabilized with pluronic acid (0.02%). Neurons were loaded with Fura2-AM, stimulated with KCl 50 mM prior addition of fCCP to reveal the mitochondrial stocks of calcium. **E**, **F**) Quantification of calcium increased upon depolarization (**E,** ratio of the fluorescence peak after adding KCl to the mean of the 10 first basal measurement) and the amount of mitochondrial calcium released by fCCP (**F,** ratio of the fluorescence peak after adding fCCP to the lowest point during wash) in normal, MPC-deficient neurons and neurons+RU360. N>13 neurons per condition from 3 independent experiments. One-way ANOVA+Holm Sidak’s post-hoc test ((**E**) Ctrl vs Zaprinast p=0.0389, Ctrl vs UK-5099 p=0.0054, Ctrl vs Rosiglitazone p=0.0001, Ctrl vs RU360 p=0.0001; (**F**) Ctrl vs all conditions p=0.0001). **G, H)** Example traces and F-I relationship in WT cells with standard intracellular solution and with a solution containing the MCU inhibitor RU360 (1 or 10 μM), which increased neuronal firing (10 μM: Two-way ANOVA, F(1, 72) = 26.03, p<0.0001). **I, J)** Lack of RU360 (10 μM) effect on neuronal firing in KO cells (Two-way ANOVA, F(1, 72) = 0.03607, p = 0.8499). **K)** Example traces of WT cell firing elicited by a current ramp (300 pA max amplitude, APs are trimmed) in control condition and with RU360, with expanded portion at the bottom indicating mAHP measurement. **L)** Summary graph of mAHP values in control condition and with RU360, indicating significant reduction in WT (unpaired t test, t = 2.352, p = 0.0392). **M, N)** Example traces and F-I relationship in WT cells infused with RU360 (10 μM) and subsequently exposed to βHB (2 mM, >20 min exposure), which decreased neuronal firing (Two-way ANOVA, F(1, 28) = 17.69, p = 0.0001) and augmented mAHP (inset, paired t test, t = 2.336, p = 0.0477).

Mitochondria import calcium through the mitochondrial calcium uniporter (MCU) in a membrane potential dependent manner and thereby play a major role in calcium homeostasis(*37*). In an interesting study published during the preparation of this manuscript, it was reported that inhibition of the MPC in cardiomyocytes and hepatocytes can result in higher expression of the MCU gatekeeper MICU1 and inhibition of MCU-mediated calcium uptake(*38*). However, our investigations did not validate this hypothesis (Supplementary figure 7). Therefore, we focused our study on the mitochondrial membrane potential, which we found reduced in the MPC-deficient neurons (Figure 1c, f), and which could affect the buffering capacity of mitochondria upon stimulation. This hypothesis was tested using Fura2-loaded cultured cortical neurons at rest or upon stimulation with KCl, in the presence or absence of chemical inhibitors of the MPC. These experiments confirmed that the peak and duration of calcium concentration in the cytosol were significantly increased in MPC-deficient neurons (Figure 6d-f). The mitochondrial uncoupler fCCP was then added to the cultures to release the calcium retained in the mitochondria. The amount of calcium released from MPC-deficient mitochondria by fCCP was significantly lower than in wild type controls (Figure 6d-f), suggesting that the capacity of mitochondria to import and store calcium was decreased in these neurons. Moreover, the elevation of cytosolic calcium seen in MPC-deficient neurons was recapitulated in WT neurons following the addition of RU360, an inhibitor of the MCU (Figure 6d-f). To further test whether the increased cytosolic calcium resulting from dysfunctional mitochondria was responsible for the hyperexcitability of MPC1-deficient neurons, we performed electrophysiological recordings in CA1 pyramidal cells in presence of RU360 into the patch pipette. In neuro-MPC1-WT cells, addition of 1 μM or 10 μM RU360 to the cell pipette caused an increase in cell firing (Figure 6g, h) while 10 μM RU360 had no effect in neuro-MPC1-KO cells (Figure 6i, j). Importantly, blockade of the MCU with RU360 in neuro-MPC1-WT cells was accompanied by a significant reduction in the mAHP (Figure 6k, l), indicating that calcium alterations induced by mitochondrial dysfunction may indeed affect M-type K^+^ channel activation. Finally, whereas βHB treatment did not significantly alter the firing of neuro-MPC1-WT cells in control conditions (Figure 5e, f), it reduced the excitability of cells infused with 10 μM RU360 while slightly increasing mAHP (Figure 6 m, n), consistent with the hypothesis that βHB normalizes the alteration in the M-type K+ conductance.

Altogether, our results show that MPC1-deficient neurons display a lower mitochondrial calcium buffering capacity which may explain the hypoactivity of the M-type K^+^ channel and the intrinsic hyperexcitability of neurons.

## Discussion

Tight coupling of neuronal activity to energy metabolism is essential for normal brain function. Here, to assess the contribution of metabolism in neuronal activity, we inactivated the MPC specifically in adult CamKIIα-expressing neurons in the mouse. As previously reported(*18*)(*39*), we found that loss of the MPC led to decreased OXPHOS in glutamatergic neurons. Despite this, these mice appeared normal at rest and presented a normal behavioral repertoire (i.e., novelty exploration, sociability, stress coping), except for lower anxiety-like behaviors which are consistent with a higher glutamatergic tone(*40*). However, they developed severe seizures immediately following low level administration of two pro-convulsant drugs, the GABA receptor antagonist pentylenetetrazole (PTZ), or the glutamate receptor agonist kainic acid.

The lack of an apparent phenotype in neuro-MPC1-KO mice under resting conditions suggests that, up to a certain point, mitochondria can compensate for the deficit in mitochondrial pyruvate import by using other substrates to fuel the TCA cycle. Recently, Timper et al.(*41*) reported that proopiomelanocortin (POMC)-expressing neurons, in which OXPHOS was impaired either by partial inactivation of Apoptosis-Induced Factor or by deletion of MPC1, were able to rewire their metabolism towards mitochondrial fatty acid oxidation to stimulate mitochondrial respiration. However, it seems unlikely that such a compensatory mechanism can occur in MPC1-deficient glutamatergic neurons since these neurons do not express the enzymes necessary for β-oxidation of fatty acids(*42*). Furthermore, it is unlikely that the astrocyte-neuron-shuttle, which supplies astrocyte-derived lactate to neurons to boost OXPHOS(*43*), can circumvent the loss of the MPC since all available data thus far indicate that lactate must first be converted by neuronal LDH into pyruvate in order to fuel the TCA cycle.

Instead, our data point toward aerobic glycolysis as a likely compensatory mechanism for the OXPHOS deficit as several studies have reported that increased glycolysis at the synapse, uncoupled from oxidative phosphorylation, could provide sufficient ATP to ensure normal neurotransmission(*6, 44*). It is therefore possible that, under resting conditions, increased aerobic glycolysis participates in the activity of MPC1-deficient glutamatergic neurons.

Despite the lack of an obvious phenotype in resting mice, we found that, when challenged with the pro-convulsant molecules PTZ or kainic acid, the neuro-MPC1-KO mice were far more sensitive than WT animals and rapidly exhibited severe acute seizures. This suggests that the basal electrical activity of MPC1-deficient neurons may be continuously counterbalanced by inhibitory synapses, providing the normal resting phenotype described above. However, upon release of the ‘brakes’ exerted by the inhibitory system, the neuro-MPC1-KO neurons would become hyperactive, which would translate into the observed epileptic output. Consistent with our data, mice deficient in pyruvate dehydrogenase (PDH), the enzyme acting immediately downstream of the MPC, were found to display an epileptiform cortical activity accompanied by behaviorally observable seizures(*45*). In this case, the epileptiform activity occurred in the context of reduced background cortical activation and, as suggested by the authors, the most likely explanation was that seizures resulted from a combination of decreased activity of inhibitory neurons, mostly parvalbumin-expressing cortical neurons, with slightly overexcitable excitatory neurons. Similar to PDH-deficient neurons, we found that the MPC1-deficient neurons displayed higher input resistance and increased spike frequency after stimulation, a phenotype that we investigated further and found to be mediated by an impairment of the medium component of the after-hyperpolarization potential mediated by an M-type K+ conductance.

K^+^ efflux is the primary force behind the cellular repolarization that limits the spike after depolarization and thereby prevents neuronal hyperexcitability. One important class of K^+^ channels that fulfills this task is the M-current (*I*M)-generating KCNQ channel family (also called Kv7 channels)(*46*). In hippocampal neurons the *I*M is mediated by the KCNQ2 and KCNQ3 channels (Kv7.1 and Kv7.2), which form hetero or homodimers. Loss of function of KCNQ2 or KCNQ3 causes epilepsy in humans and mice(*27, 47–49*). In support of the notion that these channels underlie the intrinsic membrane hyperexcitability of MPC1-KO neurons, we found that inhibition of these channels using the small molecule XE991 did not change the electrical properties of KO neurons, while it made WT neurons more excitable. Our results suggest that KCNQ2/3 channels are closed in MPC1-deficient neurons, and that this could underlie their hyperexcitability. The reason for the silencing of these channels appears to be linked to an excess of cytosolic calcium. The calcium binding protein, Calmodulin (CaM), has been shown to bind to the C-terminal part of the KCNQ channel and to be required for its activity(*34*). Intracellular calcium decreases CaM-mediated KCNQ channel activity(*32, 36)* by detaching CaM from the channel or by inducing changes in configuration of the calmodulin-KCNQ channel complex(*36*). Importantly, increasing cytosolic calcium levels in wild type neurons from acute hippocampal slices, using the MCU inhibitor RU360, was sufficient to increase their firing properties, while RU360 had no significant effect on the excitability of the neuro-MPC1-KO neurons. The increased intracellular levels of calcium in neuro-MPC1-KO neurons probably result from a decreased capacity of mitochondria to buffer cytosolic calcium. Such a reduction in calcium buffering is likely to be the consequence of a reduced mitochondrial membrane potential, directly linked to a decreased oxygen consumption and OXPHOS. It is known that cells with low respiratory capacity consume ATP through the ATP synthase to maintain a minimal mitochondrial membrane potential that allows them to survive(*50*). Accordingly, ketone bodies, which restore oxygen consumption, ATP production, and mitochondrial membrane potential, reduce the excitability of neuro-MPC1-KO neurons. In addition our study suggest that βHB could act directly on the M-channel, as previously reported by Manville et al.,(*31*).

In conclusion, using mice carrying an inducible deletion of the MPC specifically in excitatory neurons, we have shown that, despite impaired pyruvate-mediated OXPHOS, glutamatergic neurons can sustain high firing and trigger severe behaviourally observable seizures when the GABAergic network is inhibited. Furthermore, our data provide an explanation for the paradoxical hyperactivity of excitatory neurons resulting from OXPHOS deficits, which often accompanies neuropathologies such as cerebral ischemia or diverse mitochondriopathies, and identify KCNQ channels as interesting therapeutic targets to prevent seizures occurring in these pathologies.

## Material and Methods

### Study design

Data sources from mice included in vivo (behavioural tests, pro-convulsant drug injections, electroencephalogram), brain slice recordings of neuronal activity and electrophysiology, isolation of synaptosomes and primary culture of cortical neurons. For mouse experiments, pilot data from three or four samples per group provided an estimate of SD and effect magnitude, which, together with a power of 0.8 and P < 0.05, guided sample sizes using the G*power software (G*power version 3.1.9.6.). MPC1-WT and MPC1-KO mice from the same litter were randomly selected for experiments. Replicates and statistical tests are cited with each result. All procedures were approved by the Institutional Animal Care and Use Committee of the University of Geneva and with permission of the Geneva cantonal authorities. Data analysis was blind and performed concurrently on control and experimental data with the same parameters. No data, including outlier values, were excluded.

### Mice

The CamKIIα-CreERT2 mouse was obtained from Jackson (stock number 012362). The MPC1^Flox/Flox^ mouse was a gift from professor Eric Taylor (University of Iowa)(Gray). The Ai14 reporter mouse was a gift from professor Ivan Rodriguez (University of Geneva)(Madisen). The GFAP-CreERT2 mouse was a gift from professor Nicolas Toni (University of Lausanne)(*51*). By using the Cre driver lines, we generated two different cell-type specific MPC1-KO mice: CamKIIα-CreERT2^+^-MPC1^Flox+/Flox+^ mice (here called neuro-MPC1-KO) in which MPC1 was knocked out specifically in excitatory glutamatergic neurons; and GFAP-CreERT2^+^-MPC1^Flox+/Flox+^ mice in which MPC1 is knockout specifically is astrocytes (here called astro-MPC1-KO). In all experiments age-matched wild type controls were used and are referred to in the text as neuro-MPC1-WT (CamKIIα-CreERT2^-^-MPC1^Flox+/Flox+^) and astro-MPC1-WT mice (GFAP-CreERT2^-^-MPC1^Flox+/Flox+^). The neuro-MPC1-KO and astro-MPC1-KO phenotypes were tamoxifen-inducible. In order to induce MPC1 deletion, the mice were injected intraperitoneally (ip) for 5 consecutive days with 100μl of 10mg/ml tamoxifen (Sigma, 85256) in sunflower oil. The mice were considered to be MPC1-KO from one week after the final injection. All experiments were carried out in accordance with the Institutional Animal Care and Use Committee of the University of Geneva and with permission of the Geneva cantonal authorities (Authorization numbers GE/42/17, GE/70/15, GE/123/16, GE/86/16, GE/77/18, GE/205/17) and of the Veterinary Office Committee for Animal Experimentation of Canton Vaud (Authorization number VD3081).

### Pentylenetetrazol (PTZ)-induced convulsion protocol

We used the PTZ kindling model of epilepsy as described in Dhir et al.,(*21*). Briefly, this test entails chronic intraperitoneal (ip) injection of 35 mg/kg PTZ (Sigma, P6500), which is a sub-convulsant dose for WT mice, every 2 days for 2 weeks, and after each PTZ injection, the mice were scored according to their clinical symptoms, as described in previously (*21),(52*). After each PTZ injection, the animals were gently placed in isolated transparent plexiglass cages and their behaviour was observed to assign a seizure score based on the following criteria: stage 1: sudden behavioral arrest and/or motionless staring; stage 2: jerks; stage 3: Straub’s tail (rigid tail being held perpendicularly to the surface of the body); stage 4: partial clonus in a sitting position; stage 5: generalized clonus; stage 6: convulsions including clonic and/or tonic-clonic seizures while lying on the side and/or wild jumping (convulsive status epilepticus). Mice were scored over a period of 30 min and the tests were performed in semi-blind mode (carried out by 2 experimenters of which only one knew the genotype). After the PTZ test, mice were immediately sacrificed in a CO2 chamber. The seizure severity score was calculated by taking the sum of the behavior and seizure patterns for all animals in a group and dividing by the number of animals present in the group.

### Electroencephalogram (EEG)

Surface EEGs were recorded in head-fixed, awake animals with 32 stainless steel electrodes (500 μm Ø) covering the entire skull surface as described previously(*53, 54*). Briefly, a headpost was placed under isoflurane anaesthesia allowing head-fixation. Recording sessions took place after a period of 4 days of head-fixation training to allow acclimatization of the animals to the experimental setup. PTZ was injected ip at the beginning of the session. Electrophysiological differential recordings were acquired with a Digital Lynx SX (Neuralynx, USA) at a sampling rate of 4 kHz and with a 2kHz low-pass. The ground electrode was placed above the nasal bone and the reference electrode was placed on the midline between parietal bones (channel 31, Figure 3C). All signals were calculated against the average reference offline.

### Patch-clamp electrophysiology

Tamoxifen-treated MPC1Flox/Flox-CamKIIαCre(+) mice and wild-type littermates (6-10 weeks-old) were anaesthetized with isoflurane and decapitated, and the brain was quickly removed and placed in oxygenated (95% O2 / 5% CO2) ice-cold N-Methyl-D-glucamine (NMDG)-based medium, containing (in mM): 110 NMDG, 2.5 KCl, 1.2 NaH_2_PO_4_, 30 NaHCO_3_, 20 HEPES, 10 MgCl_2_, 0.5 CaCl_2_, 25 glucose, 5 L(+)-ascorbic acid, 2 thiourea, 3 Na-pyruvate (titrated to pH 7.2-7.3 with HCl). Acute hippocampal transverse slices (350 μm thick) were cut using a vibrating tissue slicer (Campden Instruments). Slices recovered for 1 h at 35°C and subsequently at room temperature in a storage solution containing (in mM): 92 NaCl, 2.5 KCl, 1.2 NaH_2_PO_4_, 30 NaHCO_3_, 20 HEPES, 2 MgCl_2_, 2 CaCl_2_, 25 glucose, 5 L(+)-ascorbic acid, 2 thiourea, 3 Na-pyruvate (titrated to pH 7.2-7.3 with NaOH). In the recording chamber, slices were superfused with oxygenated standard artificial cerebrospinal fluid (aCSF) containing (in mM): 130 NaCl, 25 NaHCO_3_, 2.5 KCl, 1.25 NaH_2_PO_4_, 1.2 MgCl_2_, 2 CaCl_2_, 18 glucose, 1.7 L(+)-ascorbic acid.

Whole-cell patch clamp recordings were performed at nearly physiological temperature (30-32°C), with borosilicate pipettes (3-4 MΩ) filled with (in mM): 130 KGluconate, 10 KCl, 10 HEPES, 10 phosphocreatine, 0.2 EGTA, 4 Mg-ATP, 0.2 Na-GTP (290-300 mOsm, pH 7.2-7.3). A control experimental series was conducted with narrow pipettes tips (9-10 MΩ) filled with (in mM): 130 KGluconate, 5 KCl, 10 HEPES, 5 Sucrose (275-280 mOsm, pH 7.2-7.3), in order to delay intracellular dialysis(*45*) and minimize interference with intracellular ATP and Ca^2+^ levels. In this series, neuronal firing was measured within the first 1.5 min after whole-cell establishment (supplementary figure 8).

To elicit neuronal firing, cells were held at −60 mV with direct current injections, and somatic current injections of increasing amplitude were provided using ramps of 5 s (6 ramps with final amplitude ranging from 50 pA to 300 pA) or squared pulses of 2 s (25 pA delta increase, max amplitude 200 pA). Input resistance (Ri) was assessed by the passive current response to a −10 mV hyperpolarizing step while cells were held at −60 mV. In control condition, resting membrane potential (Vrmp) and neuronal firing were measured within the first 5 min from the establishment of the whole-cell condition. The rheobase and the firing threshold were measured as the level of current and voltage, respectively, that induced the first action potential in the ramp protocol. The effect of β-hydroxybutyrate (2 mM) was assessed after >20 min perfusion, and compared to cell firing prior to perfusion.

Signals were acquired through a Digidata1550A digitizer, amplified through a Multiclamp 700B amplifier, sampled at 20 kHz and filtered at 10 kHz using Clampex10 (Molecular Devices).

### Cell culture and lentiviral transduction

Wild type pregnant mice were decapitated and E18 embryos were collected in HBSS medium. Primary cultures of cortical neurons were prepared as described previously(*55*). For MPC1 downregulation, at 7 days in vitro (DIV), neurons were treated with lentiviral particles containing shRNA targeting MPC1 for a further 7-8 days. Briefly, to prepare viral particles, Hek293T cells were transfected with packaging and envelope expressing plasmids together with PLKO.1-shRNA control (SHC016, SIGMA) or targeting MPC1 (ShMPC1_1: CCGGGCTGCCTTACAAGTATTAAATCTCGAGATTTAATACTTGTAAGGCAGCTTTTT; shMPC1_2: CCGGGCTGCCATCAATGATATGAAACTCGAGTTTCATATCATTGATGGCAGCTTTTT), and after 72 hours the culture supernatant was collected, ultracentrifugated at 100,000 g for 2 hours.

### Determination of oxygen consumption rate (OCR) and extracellular acidification rate (ECAR)

Measurement of oxygen consumption was performed using a Seahorse XF 24 extracellular flux analyzer (Seahorse Biosciences). 80’ 000 cells were seeded in XF24 cell culture microplates and grown for 16 days. Measurement of basal and stimulation-dependent oxygen consumption was carried out at 37°C in aCSF (140 mM NaCl, 5 mM KCl, 1.2 mM KH_2_PO_4_, 1.3 mM MgCl_2_, 1.8 mM CaCl_2_, 5 mM Glucose, and 15 mM Hepes, pH 7.4). Cells were infected with control shRNA or shMPC1 as decribed above or treated with MPC1 inhibitors Zaprinast(*56*), Rosiglitazone(*57*), and UK5099(*58*) at 5, 5 and 1 μM, respectively. Cells were treated as indicated in the figure legends for 30 min before performing the assay. Basal oxygen consumption was measured before injection. At the times indicated, the following compounds were injected: oligomycin (1 μM), fCCP (4 μM), Rotenone/Antimycin A (1 μM). Each measurement loop consisted of 30 sec mixing, 2 min incubation, and 3 min measurement of oxygen consumption.

Determination of the extracellular acidification was carried out under the same conditions but in the absence of HEPES. The basal acidification rate was measured before injection. At the times indicated, the following compounds were injected: oligomycin (1 μM), 2-deoxyglucose (5 mM). Each measurement loop consisted of 2 min mixing, 2 min incubation, and 3 min measurement of oxygen consumption.

### ATP measurements

ATP measurements were performed on 15-17 DIV neurons, infected with control or MPC1 shRNA as described above, or treated 30 minutes prior to performing the assay with MPC inhibitors. Neurons were washed and scraped in PBS. Neurons were centrigugated at 1000 rpm for 5 min and resuspended in 100 mL of CellTiter Glo reagent and agigated for 2 min to allow cell lysis. After 10 min incubation, luminescence was recorded.

### Calcium imaging

E18.5 primary cortical neurons were isolated and seeded onto 35mm Fluorodishes. Neurons were treated with control or MPC1 shRNA at 7DIV and used for calcium imaging at 14-17 DIV. Neurons were loaded with 5 μM FuraFF or Fura2 (F14181 and F1221, Thermo Fisher Scientific) in recording buffer (150 mM NaCl, 4.25 mM KCl, 4 mM NaHCO_3_, 1,25 mM NaH2PO4, 1.2 mM CaCl2, 10 mM D-glucose, and 10 mM HEPES at pH 7.4) with 0.02% pluronic acid, at 37 °C and 5% CO_2_ for 30 min.

After washing, the cells were imaged in recording buffer using a custom-made imaging widefield system built on an IX71 Olympus microscope equipped with a 20× water objective. A Xenon arc lamp with a monochromator was used for excitation, exciting FuraFF or Fura2 fluorescence alternately at 340 nm ± 20 nm and 380 nm ± 20 nm and collecting emitted light through a dichroic T510lpxru or a 79003-ET Fura2/TRITC (Chroma), and a band-pass filter 535/30 nm. Neurons were stimulated using 10 μM glutamate (G1626, Sigma) and 10 μM Ionomycin was added at the end of each time course experiment as a positive control. Images were acquired using a Zyla CMOS camera (Andor) every 2-5 s. The images were then analysed using ImageJ.

Briefly, Regions of Interest (ROIs) were selected and average fluorescence intensity was measured for each channel including the background fluorescence. After subtracting the background fluorescence, the ratio between 380 and 340 nm was calculated and plotted as cytosolic [Ca^2+^] levels upon stimulation. The Mean amplitude was calculated for each cell using Graphpad Prism.

### Statistical analysis

The comparison of two groups was performed using a two-sided Student’s t-test or its non parametric correspondent, the Mann-Whitney test, if normality was not granted either because not checked (n < 10) or because rejected (D’Agostino and Pearson test). The comparisons of more than two groups were made using one or two ways ANOVAs followed by post-hoc tests, described in the figure legends, to identify all the significant group differences. N indicates independent biological replicates from distinct samples. Data are all represented as scatter or aligned dot plot with centre line as mean, except for western blot quantifications, which are represented as histogram bars. The graphs with error bars indicate 1 SEM (+/-) and the significance level is denoted as usual (*p<0.05, **p<0.01, ***p<0.001). All the statistical analyses were performed using Prism7 (Graphpad version 7.0a, April 2, 2016).

## Acknowledgments

We would like express our sincere thanks to Drs. Nika Danial, Timothy Ryan, Garry Yellen and all members of the Martinou lab for helpful scientific discussion during the course of this work. We also wish to extend our special thanks to Dr. Fabien Lanté and Dr. Anita Lüthi for advices on the electrophysiology experiments, Dr. Kinsey Maundrell for help in reviewing the manuscript, to Professor Nicolas Toni (University of Lausanne) who kindly provided us the GFAP-CreERT2 mouse, to Professor Ivan Rodriguez (University of Geneva) who kindly provided us with the Ai14 reporter mouse.

## Author contributions

JCM and ADLR conceived the project. ADLR performed and/or participated in all the *in-vivo* experiments, and analysed the data. ML designed, performed and analysed all experiments involving neuronal primary cultures and synaptosomes.SA designed, performed and analysed the electrophysiology experiments, under the supervision of CS. TM and AC performed the experiments using calcium imaging. ERF designed and performed behavioural analysis, under the supervision of CS. SM performed mouse breeding and genotyping and participated in some in vivo experiments. PS performed some of the calcium imaging experiment on cultured neurones, under the supervision of MD. AK and CQ performed EEGs. ET and JRu provided the MPC1 Flox/Flox mice and advices. JMN supervised statistical analysis. JCM, ADLR, ML and SA wrote the manuscript with input from all other authors.

## Funding agencies

Swiss National Science Foundation [31003A_179421/1 /1 and The Kristian Gerhard Jebsen Foundation to J.-C.M; Requip support from the Swiss National Science Fondation for the acquisition of the Seahorse apparatus (SNF 316030_145001). Spanish Ministerio de Economía y Competitividad (MINECO, BFU2017-84490-P and RYC-2015-18545) to PM.

## Data availability

The data that are supporting the findings of this study are available from the corresponding authors upon request.

