## Supplementary materials and legends for "Paradoxical neuronal hyperexcitability in a mouse model of mitochondrial pyruvate import deficiency"

**Supplementary material and methods**

**Food and diet**

Standard diet is composed of 4.9% fat, 24% crude proteins, 29% starch (Provimi Kliba AG, standard diet, 3800). The ketogenic diet (KD) is composed of 74.4% animal fat, 9.9% crude protein, 0.7% starch (Provimi Kliba AG, Ketogenic diet XL75:XP10).

**Glycaemia and Ketonemia measurements**

Glycaemia and ketonemia were measured in blood using the GlucoMen® Lx Plus kit (Menarini diagnostics) which has a sensitivity for ketones, mainly βHB, from 0.1 mM to 8 mM. If not stated otherwise, measurements were performed using one drop of blood from the tail-veins of mice fed ad-libitum.

**Behavioural tests**

In each behavioural test, 6- 10 adult animals per group were analyzed. They were tested in two tests to evaluate their anxiety-like responses: the elevated plus maze (EPM) and open field (OF), followed by a sociability test and a test for depression-like behaviors, the forced swim test (FST), all described below. Testing took place with at least 5 days apart. The **EPM** is a maze composed of two open and two closed arms (30 × 5 × 14 cm) separated by a central platform (5 × 5 cm). The mouse is introduced in the center of the maze facing the wall of the closed arm and allowed to explore freely for 5 min. The intensity of the lights was maintained at 12 lux on the open arms, 10 in the center and 4-3 lux in the closed. After each trial, the maze was cleaned with 5-7% ethanol and dried. Video tracking of the animal's location was performed by a camera placed above the arena. The percentage of time spent in the open arms, as a measure of anxiety, was calculated with Ethovision (Noldus SA) tracking system. The **OF** is an arena (50 x 50 cm) in which the experimental mouse is placed in one corner facing the wall and left to freely explore for 10 min. Lighting was maintained at 7 lux on the center. After each trial, the arena was cleaned with 5-7% ethanol and dried. Video tracking of the animal was recorded by a camera fixed above the arena, and images were processed using the Ethovision tracking system. The percentage of time spent in the center of the open field was taken as an indicator of anxiety. In the **Social preference (SP) test**, the preference for either a social stimulus or an inanimate object is tested. The arena consists of a rectangular, three-chambered box with a center (20 x 35 x 35 cm) and two side compartments (30 x 35 x 35 cm) with sawdust covering the floor. Retractable doorways are located between the compartments to allow or prevent access to the side chambers. In both side chambers, an object (black dummy mouse) or a juvenile mouse C57BL/6J (28 ± 2 days) are located in a wirecage (10.16 cm bottom diameter, 11 cm high, bars spaced 0.5 cm apart). Light conditions are kept at 10 lux in the center. Two consecutive days prior to testing, experimental animals are habituated to the apparatus for 10 min. Juvenile mice are also habituated to the cylinders for 30 min. In the test day, each experimental mouse is located in the center compartment with both doors closed for 5 min. The doors are then opened and the mouse allowed to explore the arena for 10 min. Behavior was video-recorded and the percentage of time each mouse spent sniffing either the object or the juvenile scored with the Observer (Noldus SA) software. The **Forced swim test (FST)** involves placing each mouse in in a beaker containing 3L of tempered water (24 ± 1°C) for 6 min. Behavior is recorded with a video camera and the immobility time spent during the last 4 min quantified with the Ethovision (Noldus S.A.) tracking system.

**Echo magnetic resonance (EchoMRI)**

The body composition was measured with an echo magnetic resonance (Echo Medical System). Briefly, the mouse is weighed before scanning. To calibrate the device, the mouse weight is introduced and the animal is placed on the holder inside the scanner. The % of lean mass is calculated considering the total body weight of each animal.

**Electrophysiological field recordings**

Acute hippocampal slices were prepared as described in the main Material and Methods section. Field recordings were conducted at nearly physiological temperature (30-32°C) in the presence of the GABA_A_R blocker picrotoxin (0.1 mM). Field excitatory postsynaptic potentials (fEPSPs) were acquired in CA1 stratum radiatum through a borosilicate pipette filled with aCSF while stimulating Schaffer collaterals (0.1 ms duration) with a tungsten concentric microelectrode (World Precision Instruments). Input-output curves of fEPSPs were constructed by displaying the fEPSP initial slope as a function of the amplitude of the presynaptic volley (FV), which was increased by augmenting the stimulus intensity (50-500 µA). Two consequent stimuli were provided with an interval of 50 ms. Paired-pulse ratio was calculated by dividing the initial slope of the second response by the initial slope of the first response.

Signals were acquired through a Digidata1550A digitizer, amplified through a Multiclamp 700B amplifier sampled at 4 kHz and filtered at 1 kHz for field recordings, using Clampex10 (Molecular Devices).

**Calcium imaging of the network dynamics**

Hippocampal slice cultures prepared as previously described(*59, 60*) were infected with virus to enable expression of the genetically encoded calcium sensor GCaMP6s under the control of the human synapsin promoter (University of Pennsylvania Vector Core). Slices were immersed in an artificial cerebro-spinal fluid containing 124 mM NaCl, 1.6 mM KCl, 1.2 mM KH_2_PO_4_, 1.3 mM MgCl_2_, 2.0 mM CaCl_2_ 10 mM Glucose, and 2.0 mM ascorbic acid, pH 7.4, at room temperature (22°C) and continuously perfused and oxygenated using a peristaltic pump. Calcium transients were recorded using a Nipkow-type spinning disk confocal microscope (Olympus) coupled with single-photon laser (excitation wavelength 488). Images were acquired through a CCD camera (Visitron Systems Evolve). Slices were imaged using a 10X 0.30NA objective (Olympus) at 8.9 Hz frame rate (i.e. 112 ms per frame). Typically, imaging covered a field of 420x420 µm containing ~200 individual neurons in the CA1 area. For treatment with pentylenetetrazol (PTZ), slices were pre-incubated with 2 mM drug for 15 minutes, and, slices were imaged after addition of 50 µM carbachol (CCh), which induces the generation of activity patterns mimicking those recorded in vivo(*61*). All drugs remained present throughout the recordings.

Calcium data analyses were performed using the Matlab-based ‘Caltracer3beta’ software (Columbia University), and custom-made scripts as previously described(*60*). This enabled a) tracing of Regions Of Interests (ROIs) corresponding to the identified GCaMP6s-expressing neuronal soma, and b) the calculation of the averaged fluorescence signal from each ROIs as a function of time. In order to distinguish the neuronal calcium activity from background activity (i.e. from nearby fibres), the fluorescence within a halo of the pixels surrounding the ROI was subtracted from the fluorescence signal recorded in the ROI. From the resulting signal, the onset and offsets of the calcium events were identified. Onsets were automatically detected when fluorescence in a given slice exceeded a threshold value (based on the background noise) for a minimum duration of 1s (based on the kinetics of GCaMP6s), and offsets were defined as the half-decay time of the event. Onset and offsets were manually corrected after automatic detection, and used to estimate 1) the frequency, the amplitude, and the duration of the calcium events, and 2) the occurrence of co-activation (e.g. neuronal synchronization) exceeding random chance.

**Isolation of synaptic membranes**

Isolation of synaptic membranes (hereafter referred to as synaptosomes) from P70 neuro-MPC1-WT and neuro-MPC1-KO mice was performed according to Chassefeyre et al.(*62*) with minor modifications. Briefly, cortices from one mouse were homogenized in 2 ml ice-cold isotonic lysis buffer (0.32 M sucrose, 4 mM Hepes pH 7.4, protease inhibitors) using a teflon-glass-homogenizer (10 strokes). The homogenate was centrifuged at 1,000 x g for 10 min to remove nuclei. The post-nuclear supernatant (S1) was spun down at 4°C for 20 min at 13,800 x g, to yield crude synaptosomes (pellet P2) and crude cytosol (supernatant S2). The P2 fraction was homogenized in lysis buffer and layered onto discontinuous sucrose density gradients consisting of 3 ml each of 0.8M, 1.0M, 1.2 M sucrose in 4mM Hepes, pH 7,4 containing protease inhibitors. The gradients were centrifuged at 82,500 x g in a Beckman SW41 rotor for 120 min and the synaptosomes were collected at the 1.0-1.2 M interface, resuspended in 5 vol. lysis buffer and spun down for 20 min at 150,000 g. The pelleted synaptosomes were resuspended in the appropriate buffer for either western blotting and oxygen consumption rate as described below under separate subheadings.

**Determination of oxygen consumption rate (OCR) on synaptosome**

Purified synaptosomes were resuspended in 140 mM NaCl , 5 mM KCl, 5 mM NaHCO_3_, 1 mM MgCl_2_, 1,2 mM Na_2_HP0_4_, 10 mM D-glucose, 20 mM Hepes pH 7.4 , and plated in XF24 cell culture microplates pre-coated with poly-D-lysine (100 μg/well). The plates were centrifuged at 3400 x g for 1 hour to ensure attachment of synaptosomes. Oxygen consumption was measured as for neurons (described in the main ‘Material and Method’ section), except that at the times indicated, the injected compounds were: oligomycin (2 μM), fCCP (4 μM), Rotenone/Antimycin A (1 μM), and each measurement loop consisted of 30 sec mixing, 2 min incubation, and 2 min measurement of oxygen consumption.

**Glucose uptake**

Glucose uptake measurements were performed on 15-17 DIV neurons, infected with shRNA against MPC1 or control shRNA as described above, and treated as indicated in the figure legends 30 minutes prior to performing the assay. Neurons were incubated with 2-NDBG (2-(N-(7-Nitrobenz-2-oxa-1,3-diazol-4-yl)Amino)-2-Deoxyglucose) in aCSF (described for the cell culture) containing 2mM glucose for 15 min. After 5 min washing, live fluorescence (Ex 465nm/ Em 540 nm) was quantified using Cytation 3 plate reader (BioTek Intruments).

**Immunostaining**

Immunostaining was performed according to De la Rossa et al.(*63*) Briefly, neuro-MPC1-KO and neuro-MPC1-WT mice were perfused with 1x PBS followed by 4% paraformaldehyde (PFA) in PBS and the brains where immediately post-fixed with 4% PFA at 4°C overnight. Fixed brains were cut into 30 μm sections with a vibratome (Microm HM 650V). The primary antibody was rabbit anti-MPC1 (Sigma, HPA045119, Anti-Brp44l); the secondary antibody was goat anti-rabbit Alexa Fluor® (Life technologies, A11034) in presence of DAPI. Images were acquired using fluorescent (Zeiss Axiophot) or confocal (Zeiss LSM780) microscopy.

The TUNEL assay was performed on fixed coronal sections using the DeadEnd™ Colorimetric TUNEL System (Promega, G7130) according to the manufacturer’s instructions.

For in vitro immunostaining, 15 DIV neurons infected for 7 days with MPC1-targeting shRNA or control control shRNA, were washed in HBSS and fixed in PBS containing 4% PFA and 4% sucrose for 15 min. Neurons were incubated in 5% goat preimmune serum (GPi) diluted in 50mM Tris pH7.4, 0.2% Tween20, for 20 min, then incubated in anti-MPC1 and anti-βIII-tubulin (Mouse, Biolegend-801201) antibodies for 1 h at room temperature. After several washes in PBS, the sections were further incubated in PBS, 5% GPi, 0.2% Tween20 containing goat anti-rabbit Alexa Fluor594 (Life technologies, A11037) and goat anti-mouse Alexa Fluor488 (Life technologies, A32723) in presence of DAPI. Finally, sections were air-dried and mounted as above and images were acquired by confocal microscopy (Zeiss LSM780).

**Western blotting**

Cultured neurons, brain extracts or synaptosomes were homogenized in lysis buffer (50 mM Hepes, pH 7.4, 0.5% Triton X-100, protease inhibitors), and resolved by SDS-PAGE in 8 to 15% polyacrylamide gels. Proteins were electrotransferred to PVDF membranes. Membranes were blocked for 30 min in TBS containing 0.1% Tween20 and 5% milk powder. Primary antibodies were prepared in the same blocking solution and incubated with the membranes overnight at 4°C. After 3 washes in TBS 0.1% Tween20, secondary HRP-coupled antibodies in blocking solution were added for 60 min with agitation at room temperature. After extensive washing, bound-antibodies were revealed using the ECL kit (Biorad). Antibodies used: anti-MPC1 (Rabbit, Sigma, HPA045119), anti-MPC2 (Mouse, Millipore, MABS1914), anti-synaptophysin (Mouse, Abcam, ab8049), anti-Tyrosine Hydroxylase (Rabbit, Millipore, AB152), anti-CamKIIα (Goat, Abcam, ab87597), anti-GFAP (Mouse, Sigma, G3893), anti-VDAC (Goat, Santa Cruz, sc-8829), anti-Actin (Beta-actin-peroxidase, Sigma, a3854), anti-IgG-Rabbit-HRP (Dako, P0217), anti-IgG-Mouse-HRP (Dako, P0447, anti-IgG-Goat-HRP (Santa Cruz, sc-2304).

**Supplementary figure legends**

**Figure S1. A)** Western blot analysis of lysate from cultured astrocytes (left) and cortical neurons (right). Glial Fibrillary Acidic Protein (GFAP) was used as an astroglial marker, synaptophysin was used as a neuronal marker and actin as a loading control. This analysis shows that the cultures of cortical neurons are devoid of astrocytes. **B)** Western blotting of lysates prepared from cortical neurons expressing shCtrl or different shRNAs targeting MPC1 (shMPC1_1 and shMPC1_2). Both MPC1 and MPC2 expression is decreased in neurons expressing shMPC1. Actin was used as a loading control. **C)** Immunostaining of cortical neurons expressing shCtrl or shMPC1 for MPC1 (red) and anti-βIII-tubulin (green), a neuronal marker. Note the absence of MPC1 immunostaining in neurons expressing shMPC1 (scale bar: 20 μm) **D)** Oxygen consumption rates of cortical neurons cultured in the presence or absence Zaprinast (5 μM), Rosiglitazone (5 μM) and UK5099 (1 μM) for 1 hour, using the Seahorse XF analyzer. Assays were performed in the presence of pyruvate (5 mM) and glucose (5 mM) as carbon sources. Quantification of basal OCR expressed as a ratio to the control condition, DMSO. N>20 independent experiments. One-way ANOVA followed by Holm Sidak post hoc test (DMSO vs all conditions p=0.0001). **E)** Profile and quantification of the extracellular acidification rate (ECAR) in cortical neurons expressing shCtrl or shMPC1. Left panel: the following compounds were injected at the times indicated by vertical lines: Oligomycin (oligo.), 2-Deoxyglucose (2DG); Right panel: quantification of ECAR is expressed as a ratio to the shCtrl control condition. N=15 independent experiments. Unpaired t test (shCtrl vs shMPC1 p=0.0026). **F)** 2-NBDG uptake in neurons expressing shCtrl or shMPC1. N=15 independent experiments. Unpaired t test (shCtrl vs shMPC1 p=0.0001). **G)** Profile and quantification of the extracellular acidification rate (ECAR) in cultures of primary cultures of cortical neurons expressing shCtrl or shMPC1 +/- βHB (10 mM). Left panel: the following compounds were injected at the times indicated by vertical lines: Oligomycin (oligo.), 2-Deoxyglucose (2DG); Right panel: quantification of ECAR is expressed as a ratio to the shCtrl condition. N>4 independent experiments. One-way ANOVA followed by Holm Sidak post hoc test (shCtrl vs shMPC p=0.0001, shMPC1 vs shMPC1 + βHB p=0.0001). **H)** 2-NBDG uptake in neurons expressing shCtrl or shMPC1 +/- with βHB (10 mM) 30 min prior experiment when indicated. N>6 independent experiments. One-way ANOVA followed by Holm Sidak post hoc test (shCtrl vs shMPC p=0.0377, shMPC1 vs shMPC1 + βHB p=0.0377).

**Figure S2**.**A)** Profile and quantification of oxygen consumption rates (OCR) of synaptosomes purified from neuro-MPC1-WT and neuro-MPC1-KO mice. Data were obtained using the Seahorse XF analyzer. Assays were performed in the presence of pyruvate (5 mM) and glucose (5 mM) as carbon sources. Left panel: the following compounds were injected at the time indicated by vertical lines: Oligomycin, fCCP, Rotenone and Antimycin A. Right panel: quantification of basal and maximal OCR is expressed as a ratio of the WT for each condition. N=10 independent mice for each condition. One-way ANOVA followed by Holm Sidak post hoc test (neuro-MPC1-WT vs neuro-MPC1-KO p=0.0001). **B)** 2-NBDG uptake in synaptosomes purified from neuro-MPC1-WT and neuro-MPC1-KO mice showing an increase of glucose import in neuro-MPC1-KO synaptosome compared to WT. N=6 independent mice for each condition. Mann Whitney test (neuro-MPC1-WT vs neuro-MPC1-KO p=0.0022). **C)** Coronal sections of cortex and hippocampus P70 of neuro-MPC1-WT and neuro-MPC1-KO mice stained for apoptotic cells using TUNEL assay (TUNEL positive cells are indicated with a red arrowhead)(scale bar: 500 μm). **D)** Quantification of total cell number in cortical L2/3 from neuro-MPC1-WT mice and neuro-MPC1-KO. N=3 independent mice for each condition. Mann Whitney test (neuro-MPC1-WT vs neuro-MPC1-KO p=0.2061). **E**-**J)** Effect of the inducible MPC1-depletion in body weight (**E)**, lean mass percentage (**F**), elevated plus maze (EPM) (**G**), open field (OF) (**H**), social preference (SP) (**I**) and forced swim test (FST) (**J**). N=6-10 animals/group. Unpaired t test, ((**E**,**F**,**I**,**J**) neuro-MPC1-WT vs neuro-MPC1-KO p>0.9; (**G**) neuro-MPC1-WT vs neuro-MPC1-KO p=0.054; (**H**) neuro-MPC1-WT vs neuro-MPC1-KO, p=0.073).

**Figure S3. A)** Survival rate of neuro-MPC1-WT and neuro-MPC1-KO mice during the PTZ kindling protocol. neuro-MPC1-KO mice died during the test between the second and fourth injection of PTZ. **B)** neuro-MPC1-KO mice are more sensitive to the effect of kainic acid compared to neuro-MPC1-WT mice. Mice were injected ip with 20 mg/kg kainic acid (KA) and scored clinically. N=4 independent mice for each condition. Mann Whitney (neuro-MPC1-WT vs neuro-MPC1-KO p=0.036). **C)** Strategy used to generate the astro-MPC1-KO mice. Tamoxifen induces the recombination of MPC1^Flox/Flox^ alleles and the expression of the reporter protein tdTomato (Ai14) specifically in the GFAP positive cells (1. Glutamatergic neuron; 2. Astrocytes; 3. Inhibitory neuron). **D)** Coronal sections of astro-MPC1-KO-Ai14 of the cortex (upper panel) and hippocampus (lower panel) showing the expression of tdTomato in the astrocytes. L2/3, L4, L5 and L6 correspond to the respective cortical layers. CA1, CA3 and DG (dentate gyrus) correspond to the respective areas of the hippocampus. Scale bar: 100 μm or 10 μm (high mag). **E)** Western blot analysis of brain extract from astro-MPC1-WT and astro-MPC1-KO demonstrates a reduced amount of MPC1 in the KO mice whereas the neuronal marker CamKIIα is equal in WT and KO animals. N=8 independent mice for each condition. One-Way ANOVA followed Tukey’s post hoc test (astro-MPC1-WT vs astro-MPC1-KO p=0.0373). **F, G)** Astro-MPC1-KO mice behave as astro-MPC1-WT mice in the PTZ-kindling protocol. N=6 independent mice for each condition. Mann Whitney (astro-MPC1-WT vs astro-MPC1-KO p=0.999).

**Figure S4. A-B)** Glycemia **(A)** and ketonemia **(B)** in neuro-MPC1-WT and neuro-MPC1-KO mice maintained on the ketogenic diet (KD) or on a standard diet (SD). N=4 independent mice for each condition. Two-way ANOVA ((**A**) F(9,24)=6.629, p=0.0001, (**B**) F(9,24)=9.390, p=0.0001).  **C-D**) Levels of βHB in the blood from neuro-MPC1-KO mice 15 min following intraperitoneal injection of 1g/kg of βHB (**C**) or overnight fasting (**D**). N=3 independent mice for each condition. Mann Withney ((**C**) Vehicle vs 1g/kg βHB p=0.0593; (**D**) control vs O.N. fasting p= p=0.0593). **E**) Levels of βHB in the blood from neuro-MPC1-KO mice following addition of βHB 1% in the drinking water for 7 days. N=3 independent mice for each condition. One-way ANOVA followed by Holm Sidak’s post hoc test (Water vs βHB 1%, p=0.0309 for 2h and 4h).

**Figure S5. A)** Example voltage responses elicited in CA1 pyramidal cells from neuro-MPC1-WT (WT) and neuro-MPC1-KO (KO) by injection of current steps (protocol at the bottom, only four stimulations are shown). **B)** F-I relationship of action potential discharges evoked by current steps, indicating higher spiking frequency in KO cells (Two-way ANOVA, F(1, 234) = 61.77, p < 0.0001). **C)** Input resistance (R_i_) (Mann Whitney test, U = 244, p = 0.0247). **D)** Membrane capacitance (C_m_) (unpaired t test, t = 0.1329, p = 0.895). **E)** Resting membrane potential (V_rmp_) (Mann Whitney test, U = 302, p = 0.098). **F)** Voltage response to a subthreshold current injection of 25 pA (depol_sub_), confirming higher apparent R_i_ in KO cells (Mann Whitney test, U = 260, p = 0.0306). **G)** Rheobase, calculated as minimal current needed to elicit firing with ramp injections (Mann Whitney test, U = 211, p = 0.0046). **H)** Firing threshold (V_thres_) (Mann Whitney test, U = 252.5, p = 0.0354). **I)** HCN-mediated sag, measured as ΔV between initial and steady state voltage response to a hyperpolarizing current injection of -50 pA (Mann Whitney test, U = 299.5, p = 0.0899). **J)** Fast afterhyperpolarization (fAHP) (unpaired t test, t = 0.1315, p = 0.1942). **K)** Medium afterhyperpolarization (mAHP), measured as the negative peak of the voltage deflection at the offset of the depolarizing ramp (unpaired t test, t = 3.419, p = 0.0014). In order to compare mAHP after a similar discharge of action potentials, this measure was made using the ramps of maximal amplitude (300 pA), to which WT and KO responded with comparable firing frequencies (Figure 5b). Data are presented as mean ± SEM. # p < 0.1; * p < 0.05; ** p < 0.01. **L)** Example traces representing field excitatory postsynaptic potentials (fEPSP) recorded in CA1 stratum radiatum upon stimulation of the Schaffer collaterals with increased intensity, using a paired-pulse protocol (50 ms inter-stimulus interval). **M)** Input-output curves depicting fEPSPs slope as a function of the amplitude of the presynaptic volley (FV), showing no significant difference between genotypes (unpaired t test comparing slopes of input-output curves, t = 0.5785, p = 0.5729). **N)** Paired-pulse ratio of fEPSPs were no significantly different between genotypes (unpaired t test, t = 1.122, p = 0.29).

**Figure S6. A)** Fluorescence signal intensity from individual control neurons loaded with the calcium probe furaFF-AM (left) and Fura2-AM (right) stimulated with 10 μM glutamate prior the addition of ionomycin to reveal the calcium stock of the neurons. **B)** Fluorescence signal intensity from individual shMPC1-treated neurons loaded with the calcium probe furaFF-AM (left) and Fura2-AM (right) stimulated with 10 μM glutamate prior the addition of ionomycin to reveal the calcium stock of the neurons. **C)** Fluorescence signal intensity from individual shMPC1-treated neurons supplemented with 10 mM βHB loaded with the calcium probe furaFF-AM (left) and Fura2-AM (right) stimulated with 10 μM glutamate prior the addition of ionomycin to reveal the calcium stock of the neurons. **D)** Mean fluorescence signal intensity of cortical neurons loaded with Fura2-AM stimulated with 50 mM KCl (dashed black arrow) for 3 minutes and washed by returning in calcium free medium containing 5 mM KCl for 2 min until the level of calcium returns to basal level in the shCtrl condition.

**Figure S7.** Western blotting of lysates prepared from cortical neurons expressing shCtrl or shMPC1 for 7 days. Band intensities were expressed as a ratio with the intensity of the corresponding protein from non-transfected neurons. Note the decrease of MPC1 intensity in shMPC1 compared to shCtrl. Actin was used as a loading control. N=3 independent experiments.

**Figure S8. A)** Example voltage responses elicited in CA1 pyramidal cells from wild-type (WT) and MPC1-CamKII-KO (KO) by injection of current ramps (protocol at the bottom, only three of six ramps are displayed). Whole-cell recordings were performed with narrow pipettes (9-10 MΩ) filled with a K-Gluconate-based solution lacking ATP and EGTA. **B)** F-I relationship of action potential discharges evoked by current ramps, indicating higher spiking frequency in KO cells (Two-way ANOVA, F(1, 204) = 29.01, p<0.0001). **C)** The rheobase was reduced in KO cells (unpaired t test, t = 3.16, p = 0.004). **d)** KO cells exhibited a more hyperpolarized threshold potential (unpaired t test, t = 2.975, p = 0.0054). Data were obtained from 2 animals per genotype, and are presented as mean ± SEM. Data points represent single observations.
