## Supplementary figures and images for "Paradoxical neuronal hyperexcitability in a mouse model of mitochondrial pyruvate import deficiency"

### Figure S1

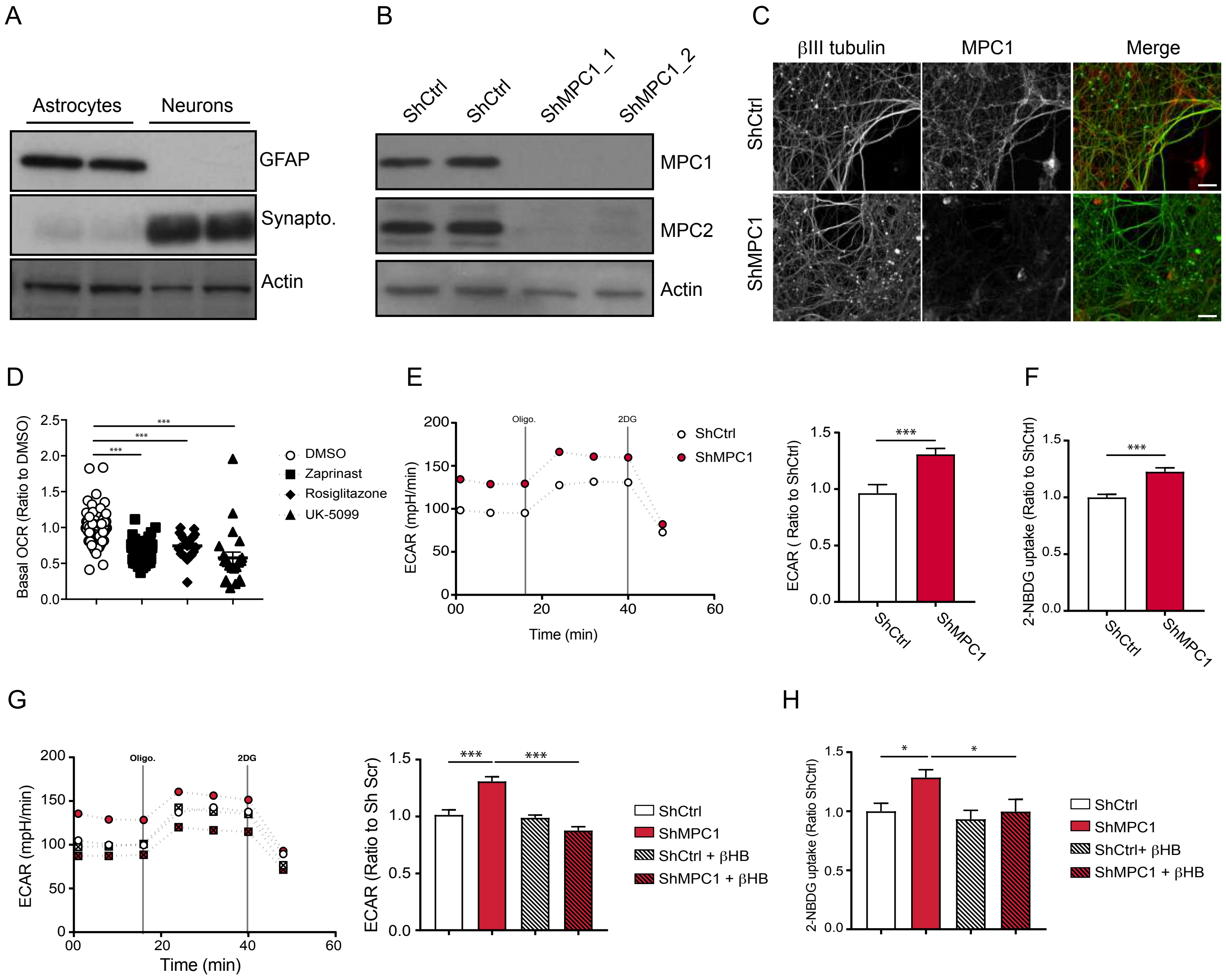

### Figure S2

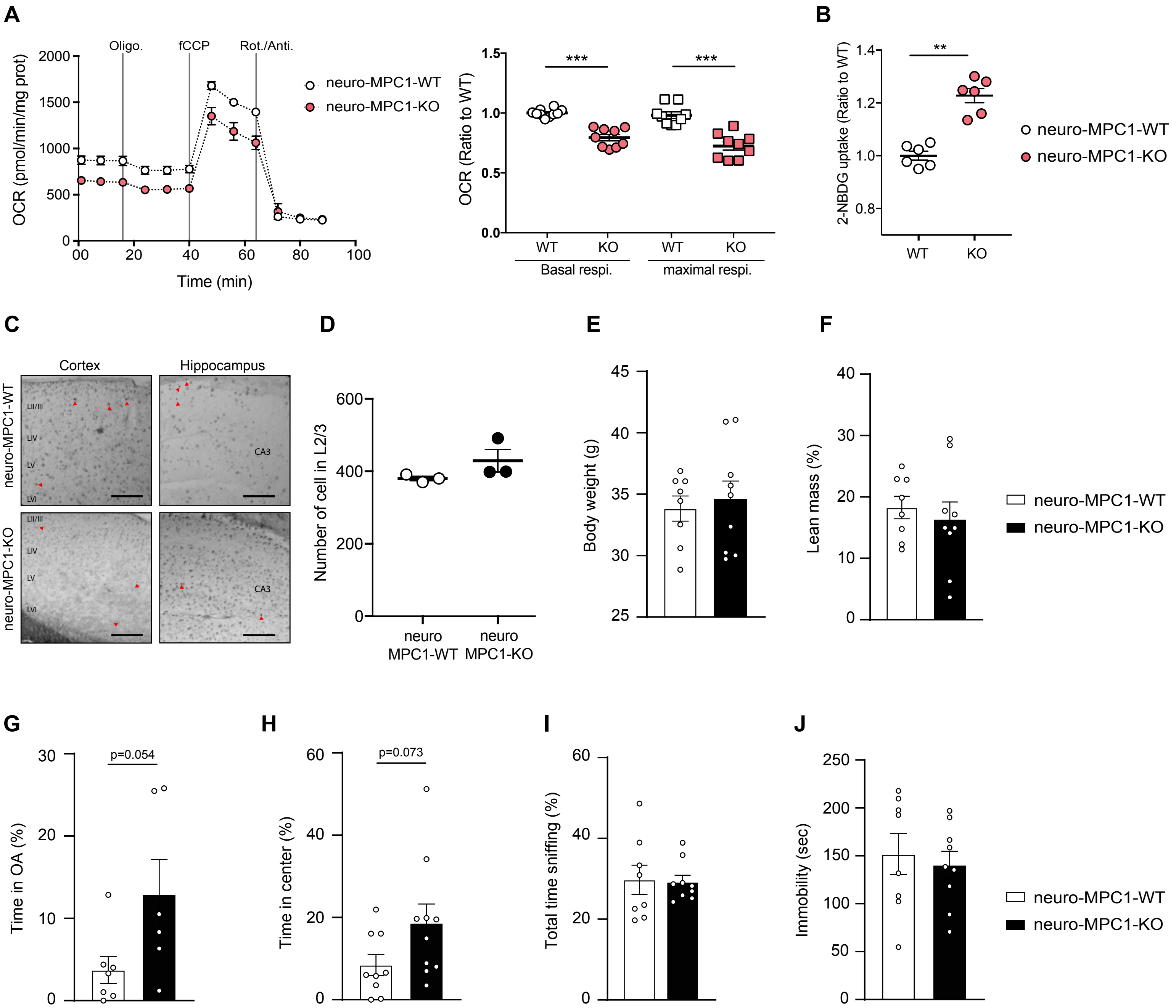

### Figure S4

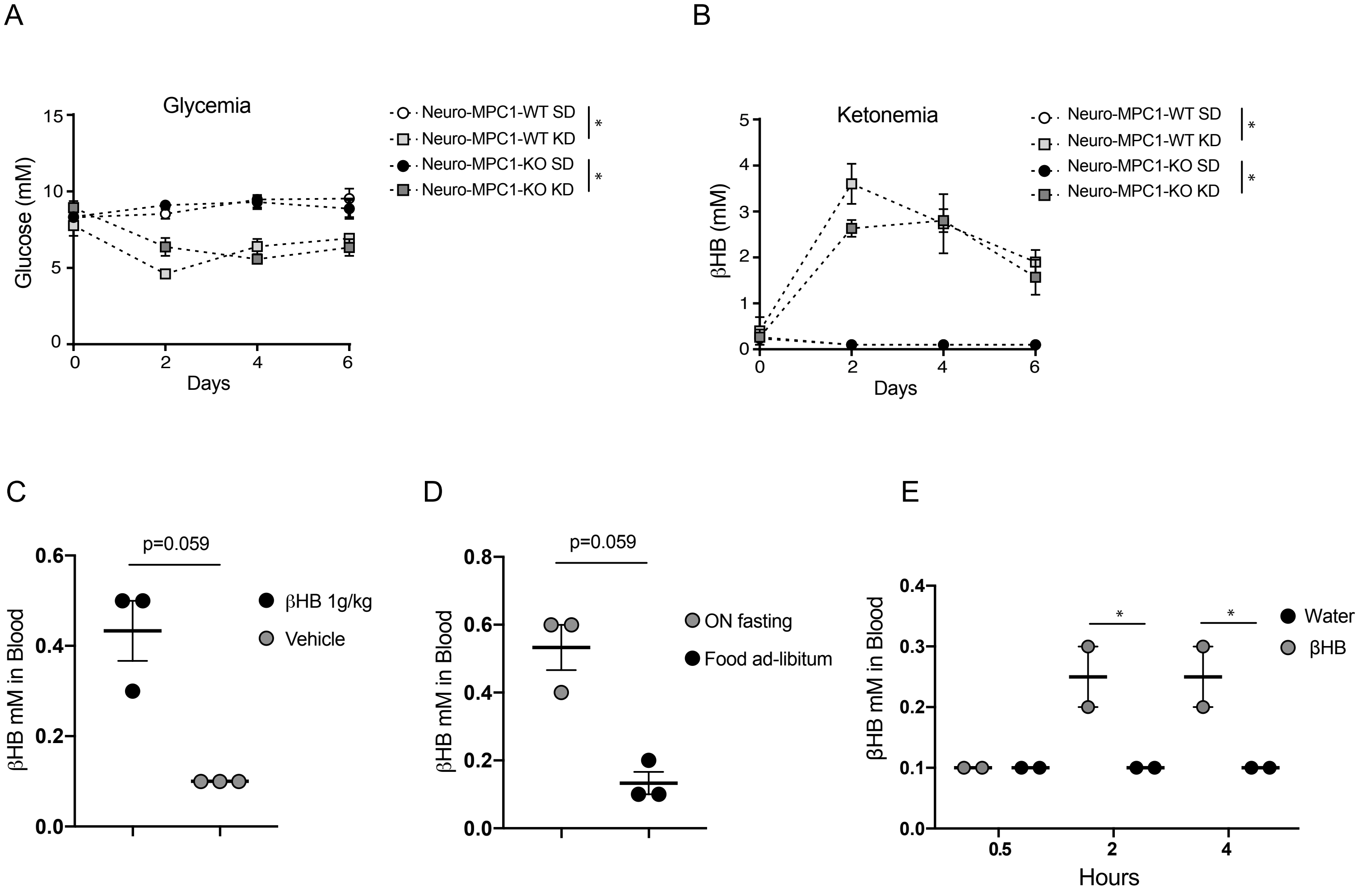

### Figure S5

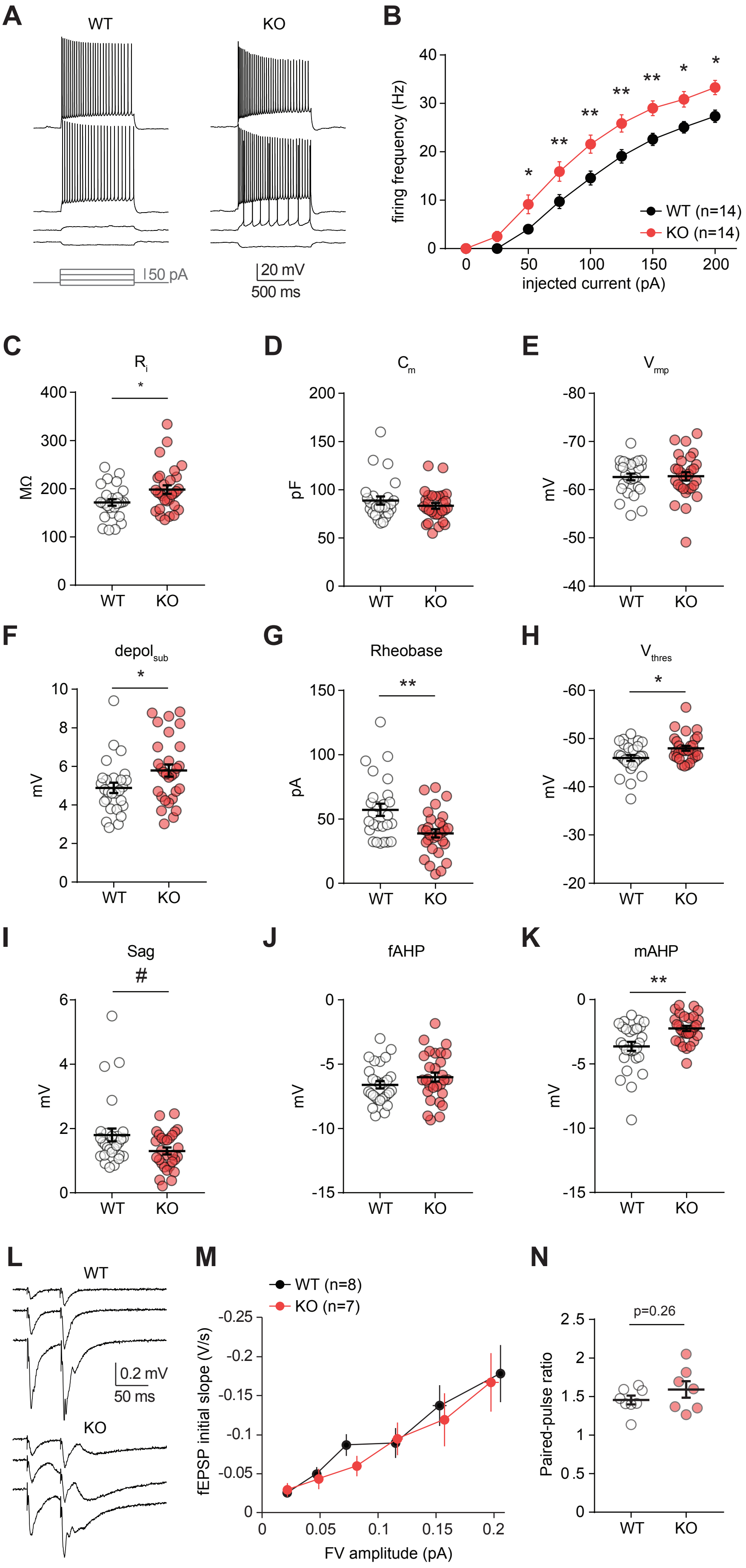

### Figure S6

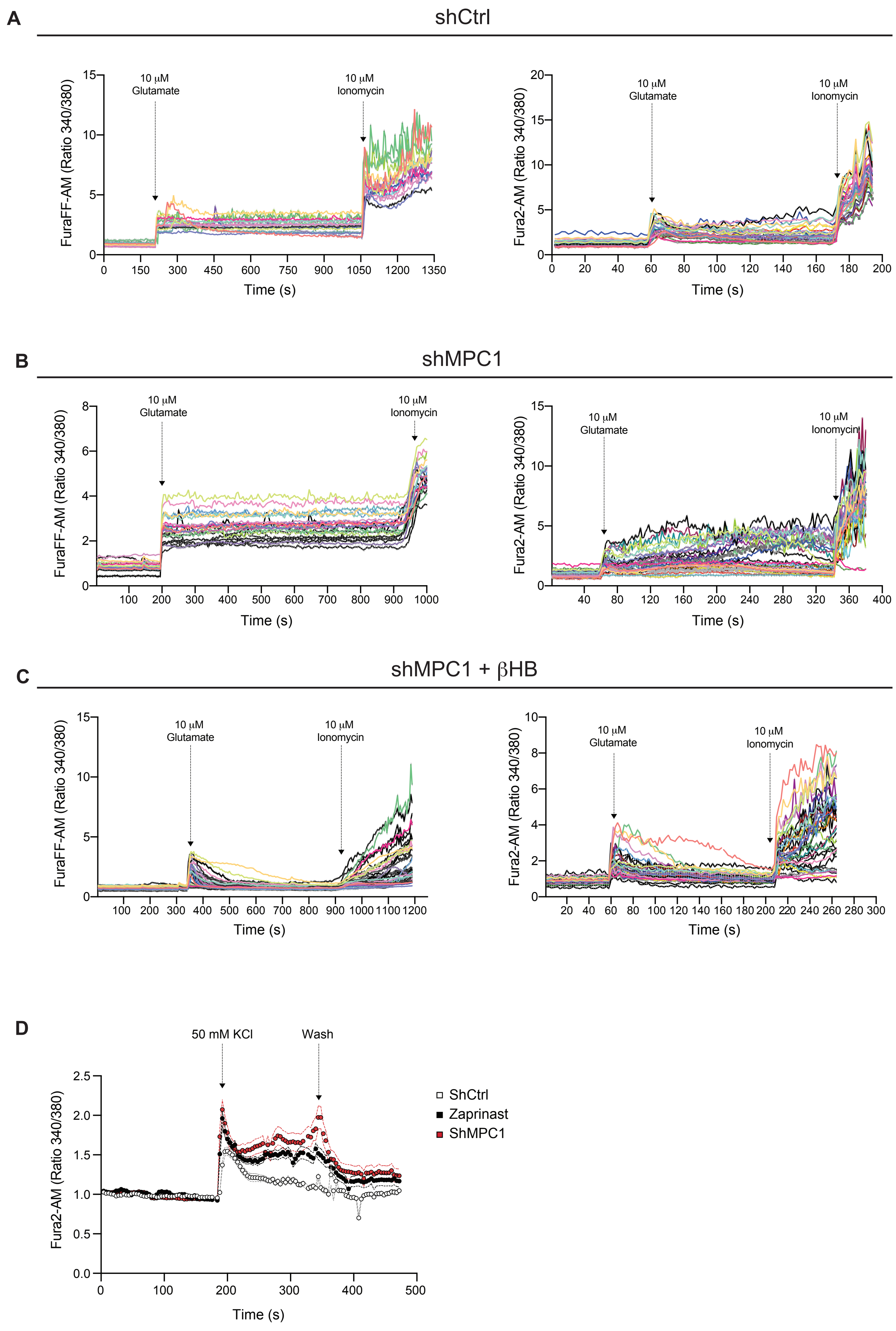

### Figure S7

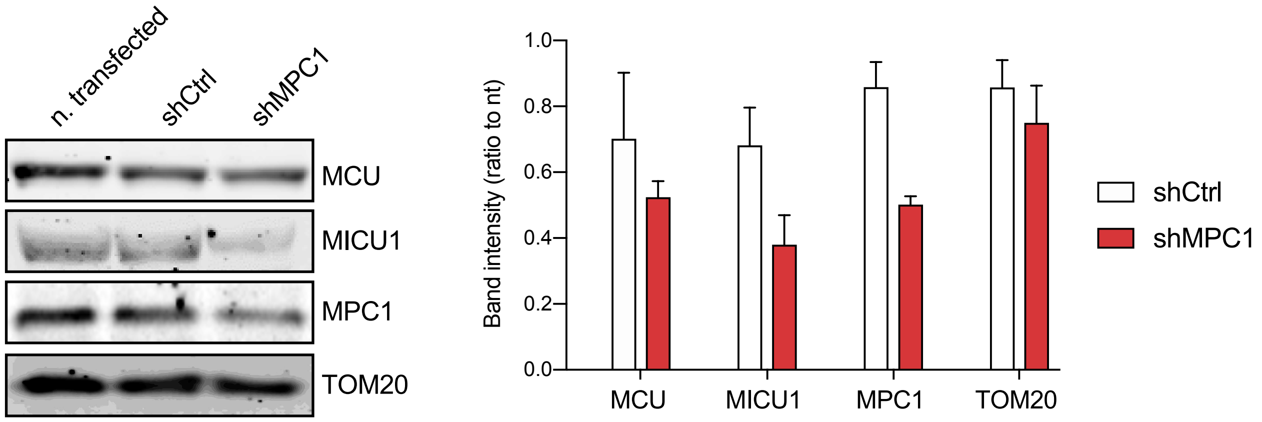

### Figure S8

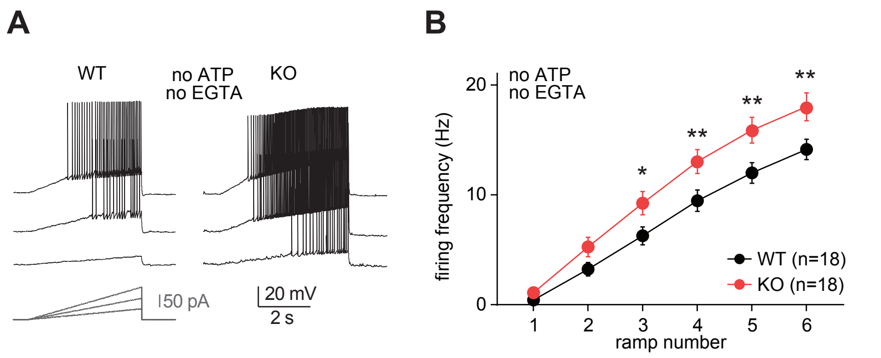
