## Supplementary material for "Paradoxical neuronal hyperexcitability in a mouse model of mitochondrial pyruvate import deficiency": Figure S3

A

*Survival rate of neuro-MPC1-WT and KO mice during PTZ-Kindling test*

| Time (days) | 1 | 3 | 5 | 7 |
|---|---|---|---|---|
| Surviving WT mice | 6 | 6 | 6 | 6 |
| Surviving KO mice | 6 | 3 | 1 | 0 |

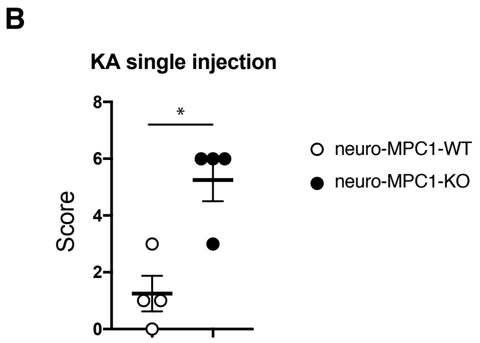


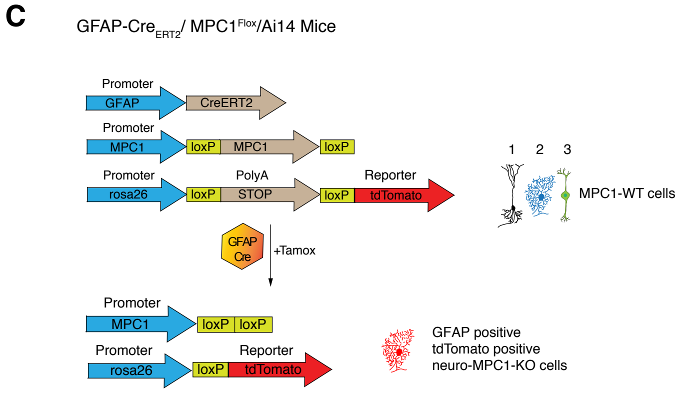


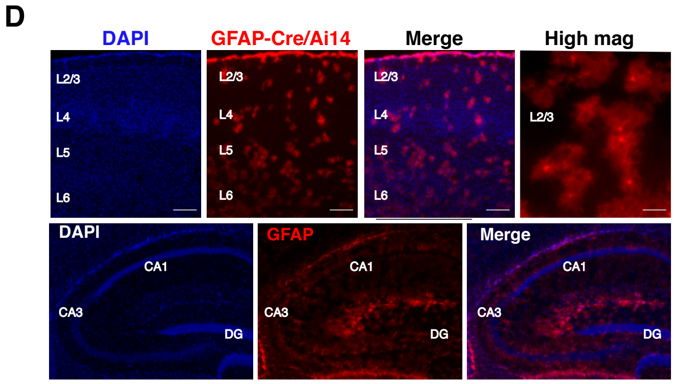


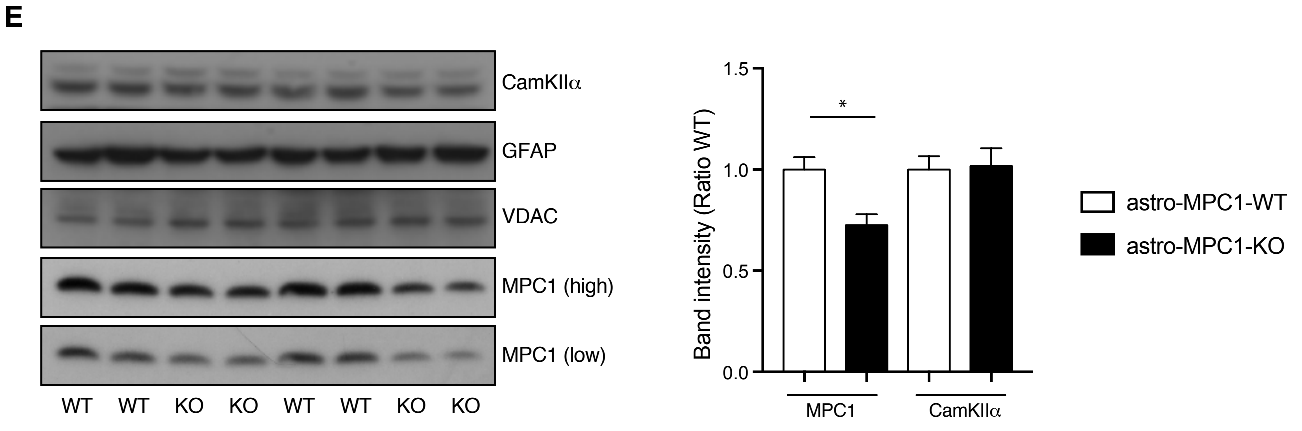


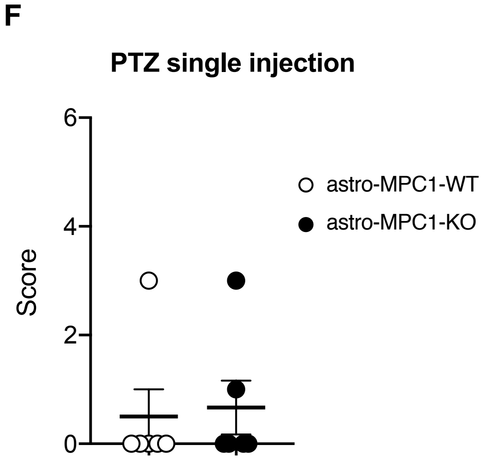


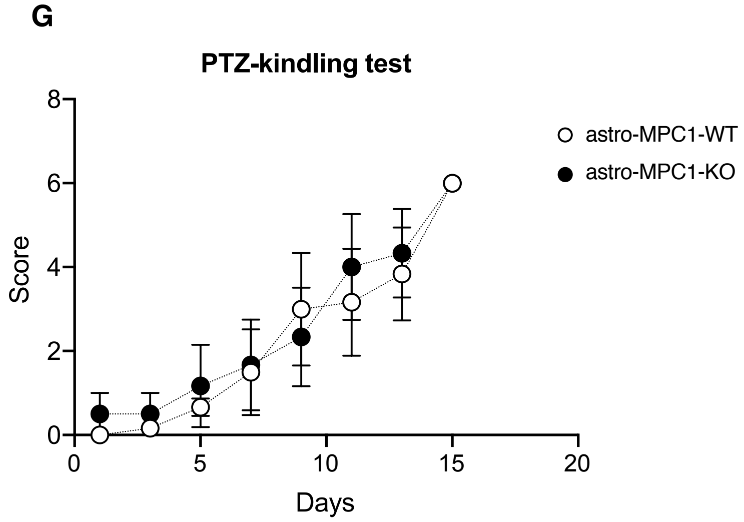


Figure S3
